# A Transcriptionally Distinct Intermediate Activation State Precedes Langerhans Cell Migration from the Epidermis

**DOI:** 10.1101/2025.05.29.656912

**Authors:** Artem Kiselev, Axel D. Schmitter-Sánchez, Sudhanshu Mishra, Hyeri Kim, Sharod Williams, Sangbum Park

## Abstract

Langerhans cells (LC) are antigen-presenting cells that form dense epidermal networks for immune surveillance. Upon antigen uptake, LCs are activated and subsequently orchestrate immune responses by migrating into the lymphatic system. However, the regulation of transcriptional programs during activation and the behavior of LCs *in vivo* during this transition remain poorly understood. Therefore, this study combines single-cell transcriptomic analysis with intravital imaging to reconstruct the activation trajectory of epidermal LCs. In particular, the study generates a high-resolution single-cell transcriptomic dataset comprising over 22,000 high-quality epidermal LCs under both homeostatic and injury conditions. Notably, a distinct intermediate activated state that precedes LC migration was identified and characterized at the level of signaling pathways and transcription factors. The integration of this dataset with external datasets from homeostatic and injured skin reveals wound-specific fibroblasts as the major source of C3, which is the central component of the complement cascade. Moreover, intravital imaging of C3-deficient mice and pegcetacoplan treatment demonstrates that C3 contributes to LC accumulation adjacent to wound sites. Collectively, the study findings identify an activated epidermal LC population and implicate complement signaling in the regulation of LC behavior during skin injury.

## 1. Introduction

Langerhans cells (LC) are a specialized subset of mononuclear phagocytes that are functionally similar to dendritic cells (DC)^[1]^. These cells function as immune sentinels by detecting environmental antigens, sensing tissue damage, and initiating immune responses, including antigen delivery to the lymph nodes^[2]^. Accordingly, LCs have been extensively studied for their development, differentiation, and immunological functions, including antigen presentation.

Additionally, LCs have emerged as key regulators of cutaneous wound healing. Specifically, LC depletion leads to excessive neutrophil infiltration, increased bacterial burden, and delayed re-epithelialization. This demonstrates their critical gatekeeper function during the early inflammatory phase of repair^[3]^. Moreover, LCs contribute to angiogenesis by associating with nascent vascular structures and activating pro-angiogenic programs^[4]^. Furthermore, LCs license effector T cells within the injured epidermis, thereby modulating immune-mediated tissue damage and repair^[5,6]^. Although LCs respond to tissue damage and inflammatory cues *in vivo*^[7–9]^, the transcriptional programs underlying LC activation and associated functional states remain poorly understood^[10–12]^.

Recent studies have revealed previously unrecognized heterogeneity and functional specializations in LC populations^[13,14]^. For example, two steady-state and two pro-inflammatory/mature LC subpopulations were identified in human skin. Notably, proinflammatory subsets are enriched in atopic dermatitis lesions, which suggests a role in disease pathogenesis^[15]^. In addition, stratified squamous epithelium contains LCs and distinct DC populations that are present in a steady state and become particularly prominent during inflammation^[10,16,17]^. These epidermal DCs overlap with populations previously described as inflammatory dendritic epidermal cells and transcriptionally and phenotypically resemble LCs^[16,18]^. Importantly, epidermal DCs express classical LC markers, including H2-E(I-E), Cd1a, and Cd207 (langerin), which renders their discrimination challenging^[16,17]^. Therefore, improved marker strategies are required to reliably distinguish LCs from other cell populations in the epidermis. This is particularly needed under inflammatory conditions. In this context, single-cell transcriptomic approaches have revealed the cellular heterogeneity and functional diversity of LCs within the epidermis. Specifically, in murine models, single-cell RNA sequencing (scRNA-seq) has shown that LCs exhibit a dual identity: functioning as tissue-resident macrophages and migrating antigen-presenting cells. This duality is influenced by intrinsic factors and the epidermal niche, thereby highlighting the complexity of the LC ontogeny and function^[19]^. Furthermore, scRNA-seq has enhanced our ability to dissect LC heterogeneity, thus enabling high-resolution profiling of individual LC populations within the epidermis^[20]^.

Accordingly, in this study, we defined the activation trajectory of LCs in response to damage-associated inflammatory stimuli using scRNA-seq and intravital imaging. Subsequently, we identified distinct LC subsets with unique gene expression signatures, thereby revealing transcriptional diversification within the injured epidermis. Overall, our findings define a previously unrecognized activation trajectory and provide a framework for understanding the mechanisms by which LCs dynamically coordinate immune responses to tissue injury.

## 2. Results

### 2.1. LCs Exhibit Similar Transcriptional Trajectories in Damage-Associated Inflammatory Models

In non-inflamed skin, LCs form a tight network across the epidermis, with the cells maintaining regular spacing^[21,22]^. This organization supports efficient immune surveillance. However, during skin insults, LCs lose their regular spatial distribution. This disorganization likely facilitates migration to draining lymph nodes to initiate adaptive immune responses^[23]^.

To define changes in LC distribution as they are activated and migrate out of the epidermis from a homeostatic state *in vivo*, we used intravital microscopy with transgenic mice expressing green fluorescent protein (GFP) under the control of the Cd207 (Langerin) promoter^[24]^. We tested whether different types of stimuli led to distinct perturbations in LC distribution (**Figure 1A**). The first method was classified as physical and involved repeated puncturing of the ear skin using a 0.2-mm microneedle array (stamp punch). The second method was classified as chemical and involved a 10-min treatment of the ear skin with a mixture of croton oil^[25]^, acetone, and olive oil (croton/acetone). Considering that fully activated LCs leave the epidermis, we aimed to identify the time point at which their spatial distribution was perturbed and they remained within the epidermis. Interestingly, in both physical damage and chemical treatments, LC spatial disorganization was most evident around 46 h post-treatment, with disrupted spatial patterns. However, the cell number was comparable with that of the homeostatic state (**Figure 1B, C**; **Video S1**). Therefore, this time point was selected for downstream analysis based on the spatial phenotype rather than the transcriptional peak of the activated state.

**Figure 1.**
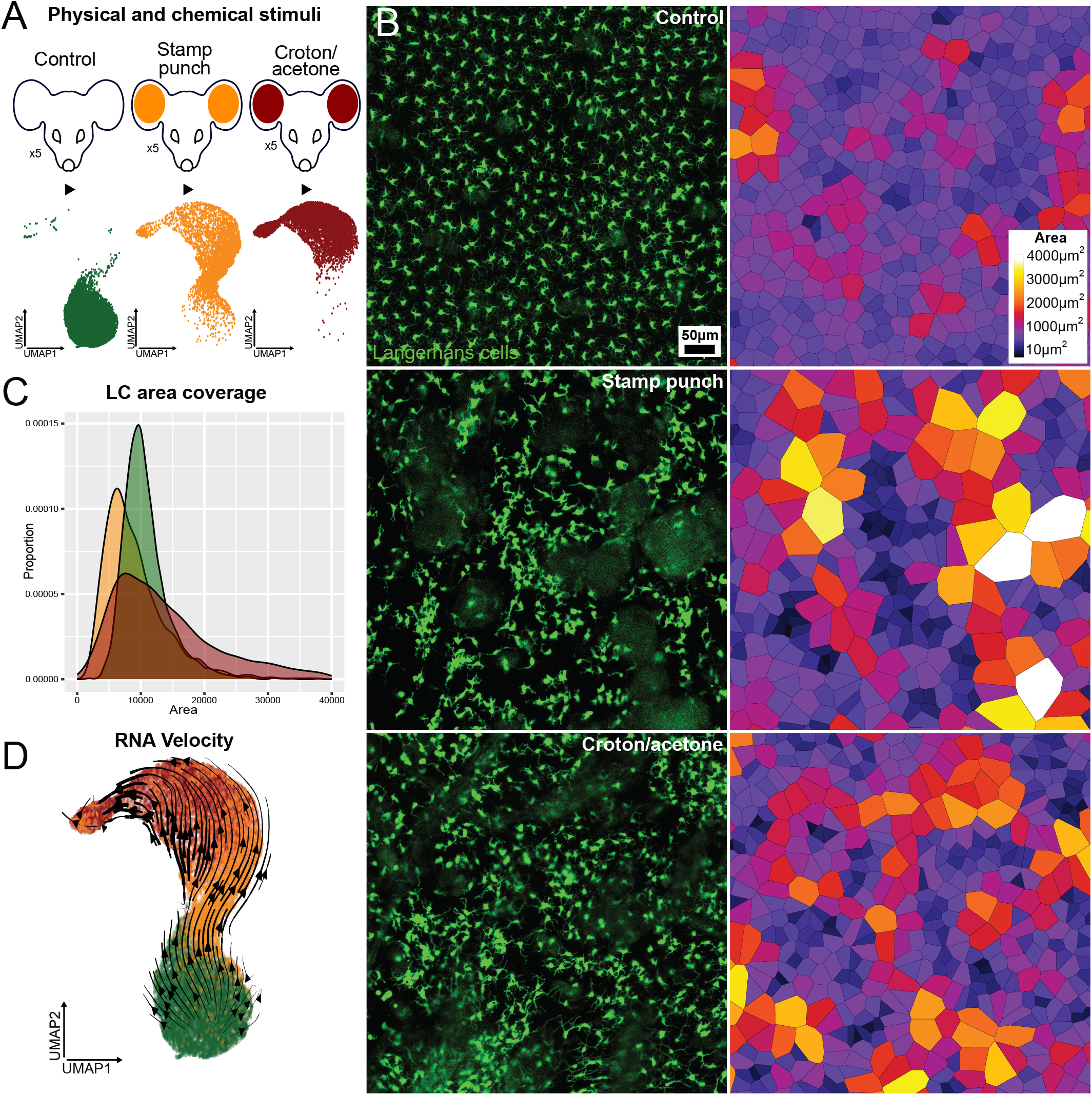
*In vivo* stimulation is associated with progressive activation and migration of LCs. **A.** Schematic overview of the experimental workflow for single-cell RNA sequencing (scRNA-seq) data acquisition. Each condition represents pooled epidermal cells from five mice. **B.** Representative intravital images of the mouse ear epidermis under three conditions: untreated control, 48 h after croton oil treatment, and 48 h after stamp-punch stimulation. Corresponding Voronoi diagrams depict the Langerhans cell (LC) spatial organization (n = 3 mice per condition). **C.** Quantification of LC territory distribution based on Voronoi diagram analysis of control and treated samples (n = 3 mice per condition). Statistical comparisons are presented in the panels. **D.** RNA velocity analysis visualized via Uniform Manifold Approximation and Projection (UMAP) embedding, with vector fields indicating inferred transcriptional directionality across the LC dataset.

Subsequently, we examined the transcriptomic changes caused by physical damage and chemical treatment via scRNA-seq using the 10x Genomics platform (**Figure S1A**). We isolated samples from three experimental groups: two corresponding to the aforementioned activation methods and one untreated control group. Each scRNA-seq sample consisted of pooled epidermal LCs extracted from the ear skin of five mice. Deep and high-quality single-cell transcriptomic profiling (**Figure S1B, C**) revealed a continuous trajectory corresponding to the progressive activation and migration of LCs, which became apparent upon the joint analysis of all three groups (treated and control; **Figure 1A**; **Figure S1D**). To validate this finding, we performed RNA velocity analysis^[26]^, which supported the directional progression from the steady state through a distinct activated state toward the migratory state in response to physical damage and chemical treatment (**Figure 1D**). Although the relative representation of these states differed between the conditions, the overall trajectory was similar across the two damage-associated inflammatory models (**Figure S2 A**–**D**).

Collectively, our intravital imaging and transcriptomic analyses demonstrate that physical and chemical stimuli induce similar activation trajectories in LCs. Across both injury models, LCs showed spatial disorganization at selected early time points, and trajectory analyses indicated progression from steady to activated toward migratory states. These results suggest that epidermal LCs engage in a shared activation program in response to damage-driven stimuli.

### 2.2. Identification of an Intermediate Activated State of LCs Preceding Epidermal Exit

Uniform Manifold Approximation and Projection (UMAP) was used to clearly define LC subpopulations during response to stimuli. Three distinct clusters were found (**Figure 2A**). Steady-state LCs representing quiescent cells in the healthy/untreated epidermis exhibit high-level expression of *Cd207*^[27]^, as well as base levels of *Cd137* (**Figure 2B, C**), *Ly6a* (**Figure 2D, E**), *Cxcr4* (**Figure 2F**), *Cd14* (**Figure 2G, H**), *Cd80*^[28]^ (**Figure 2I, J**), and *Cd86*^[29]^ markers. Gating strategies and quantification across biological replicates are shown in **Figures S3** and **S4**. These genes are associated with activation of antigen-presenting cells. The migratory state, which was identified by high expression of *Cxcr4*^[30]^ (**Figure 2F**) and *Ccr7*^[31]^, formed the smallest cluster of cells that had almost left the epidermis. This represented approximately 5% of the dataset. Between the steady state and migratory populations, we identified an intermediate activated LC population, which represented a transitional LC state that has not been well characterized^[14,32,33]^. To assess the identity of these LC populations and evaluate potential contamination from other skin cell types, we examined the established lineage markers across steady, activated, and migratory LC states (**Figure S5**). These analyses indicated that canonical LC markers were maintained across states, whereas markers of other lineages were not detectably enriched.

**Figure 2.**
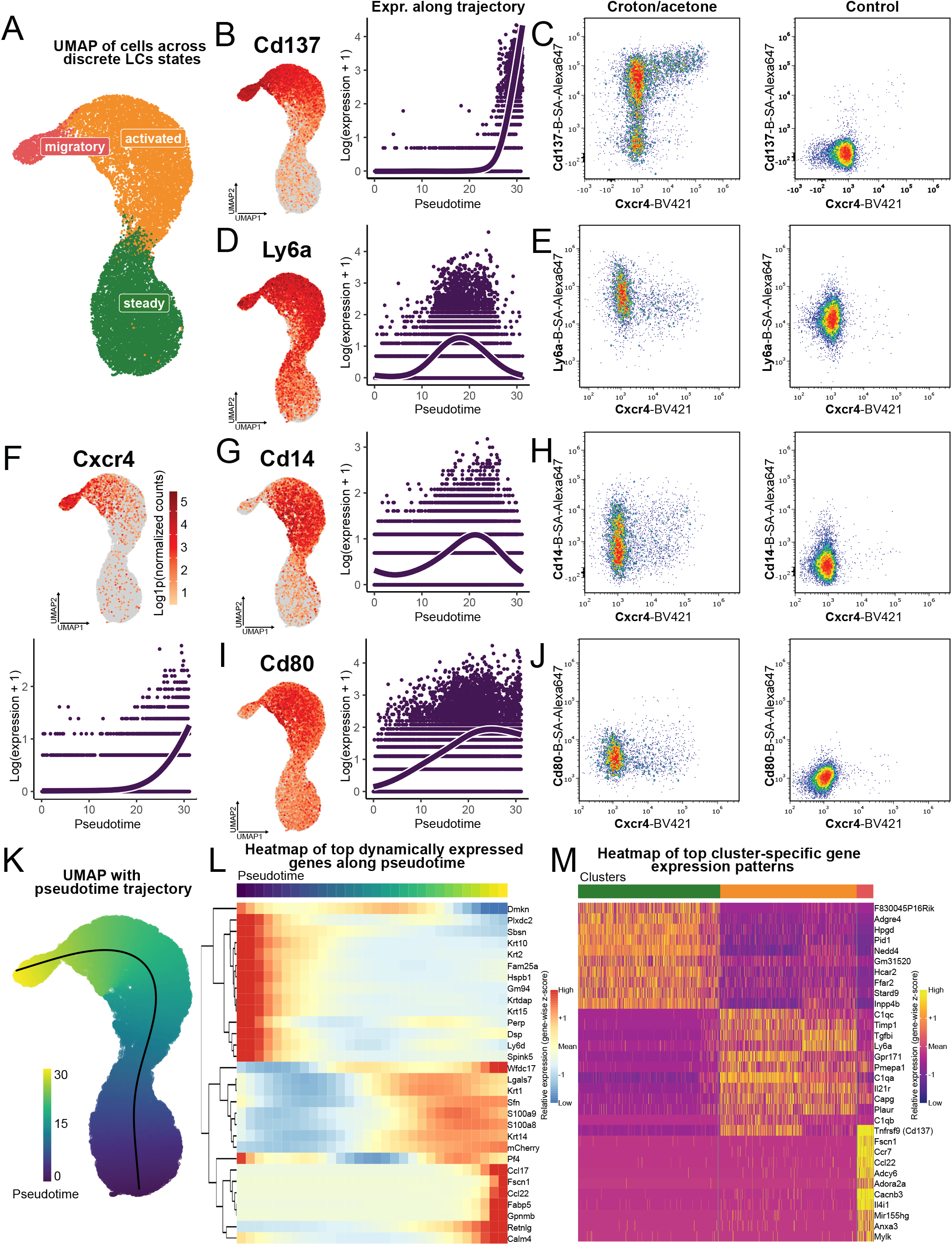
Langerhans cells (LC) transition through distinct transcriptional states along a continuous activation-to-migration trajectory. **A.** Uniform Manifold Approximation and Projection (UMAP) visualization of single-cell RNA sequencing (scRNA-seq) data showing three annotated LC clusters (steady, activated, and migratory) defined by their transcriptional profiles. **B.** FeaturePlot showing *Cd137* expression across UMAP embedding (left) and *Cd137* expression along the activation trajectory (right). **C.** Flow cytometric analysis of *Cd137* expression in LCs 48 h after croton oil/acetone treatment (left: cells pooled from three mice) and in untreated control mice (right: cells pooled from three mice). FeaturePlot showing *Ly6a* expression across UMAP embedding (left) and *Ly6a* expression along the activation trajectory (right). **E.** Flow cytometric analysis of *Ly6a* expression in LCs 48 h after croton oil/acetone treatment (left: cells pooled from three mice) and in untreated control mice (right: cells pooled from three mice). FeaturePlot showing *Cxcr4* expression across the UMAP embedding (top) and *Cxcr4* expression along the activation trajectory (bottom). FeaturePlot showing *Cd14* expression across the UMAP embedding (left) and *Cd14* expression along the activation trajectory (right). **H.** Flow cytometric analysis of *Cd14* expression in LCs 48 h after croton oil/acetone treatment (left: cells pooled from three mice) and in untreated control mice (right: cells pooled from three mice). FeaturePlot showing *Cd80* expression across the UMAP embedding (left) and *Cd80* expression along the activation trajectory (right). **J.** Flow cytometric analysis of *Cd80* expression in LCs 48 h after croton oil/acetone treatment (left: cells pooled from three mice) and in untreated control mice (right: cells pooled from three mice). UMAP visualization with pseudotime values inferred using Slingshot and tradeSeq. **L.** Heatmap showing top genes dynamically expressed along the trajectory. An extended version of this plot is shown in **Figure S6**. **M.** Heatmap of the top differentially expressed genes across the three LC clusters. An extended version of this plot is shown in **Figure S7**.

Thereafter, we aimed to further characterize the LC populations by investigating specific markers for each state, with a particular focus on the activated population (**Figure S6**, **Figure S7**). We combined bioinformatics analysis with flow cytometry-based protein-level validation. Moreover, we used differential expression analysis to identify unique markers of each LC population and shared features among groups. Our analysis identified approximately 400 transcripts that clearly distinguished activated-state LCs from steady and/or migratory states (**Table S1**). Among these transcripts, we observed upregulation of well-established antigen-presenting cell activation markers, such as *Cd80* (**Figure 2I, J**) and *Cd86* (**Figure S4**), during the activated and migratory states. In addition, during the transition to the activation and migration states, LC proliferation activity decreased, and most LCs entered the G1 stage^[34]^ (**Figure S8**). However, analysis of both transcriptomic data and flow cytometry results showed that these markers did not clearly discriminate the activated and migratory states from the steady state. To address this limitation, we prioritized the identification of alternative markers that more precisely delineate the transitions between the steady, activated, and migratory states at the transcriptomic and protein levels. In terms of the transcripts specific to the activated state, 12 candidate genes were shortlisted based on their predicted cell surface localization or roles as transcription factors. These transcripts included *Cd14*^[35]^, *Ly6a*^[36]^, *Il1b*, *Cd137* (*Tnfrsf9*)^[37]^, Ccr*7* (*Cd197), Cd49d* (*Itga4*), *Cd274*^[38,39]^, *Cxcr4*, *Adgre4* (*Emr4*), *Mafb*^[40]^, and *Bhlhe40*^[41]^. We focused specifically on transcripts with low or undetectable expression in one or more states while prioritizing those that showed a progressive increase from the steady to the migratory state or were selectively expressed in a single state with minimal expression in others. To validate these findings at the protein level, flow cytometry was performed using specific antibodies (**Figure 2C, E, H, J**). We achieved the most pronounced separation of the LC populations using a combination of surface markers. These included the new aforementioned LC activation markers (*Cd137*, *Ly6a*, and *Cd14*) and the known migratory marker *Cxcr4*. Notably, the combination of biotinylated *Cd137* and fluorochrome-conjugated *Cxcr4* provided the clearest discrimination of the activated-state LC population (**Figure 2A**). In this setup, *Cd137*-negative cells corresponded with steady-state LCs, whereas *Cd137*^+^ cells represented activated LCs transitioning toward the migratory state (**Figure 2B, C**). Additional markers, such as *Cd14* (**Figure 2G, H**) and *Ly6a* (**Figure 2D, E**), also exhibited state-specific expression patterns. In particular, expression peaked in the activated state and subsequently declined as the cells advanced toward migration (**Figure 2K–M**). To further examine the spatial distribution of activated LCs *in vivo*, we performed whole-mount staining two days after croton oil treatment or punch biopsy. CD207^+^ CD137^+^ activated LCs increased following croton oil treatment and were preferentially localized around the wound margin after punch biopsy (**Figure S9 A**–**D**).

Collectively, our findings identified an intermediate activation state of LCs between steady and migratory states. Integrated transcriptomic and flow cytometric analyses identified candidate markers associated with this transient population. This framework facilitates the identification of this state and advances our understanding of LC activation dynamics.

### 2.3. Transcriptional Reprogramming During LC Activation

To characterize the regulatory programs associated with the transitions between the LC states, the three states were subdivided into eight transcriptionally distinct substates (**Figure S1D**). Thereafter, transcription factor network activity was analyzed using SCENIC, and regulatory signaling dynamics were assessed through single-cell pathway analysis (SCPA). This enables us to track coordinated pathway changes across the LC activation continuum. Both approaches clarified the differences between the steady, activated, and migratory states (**Figure 3A–C**).

**Figure 3.**
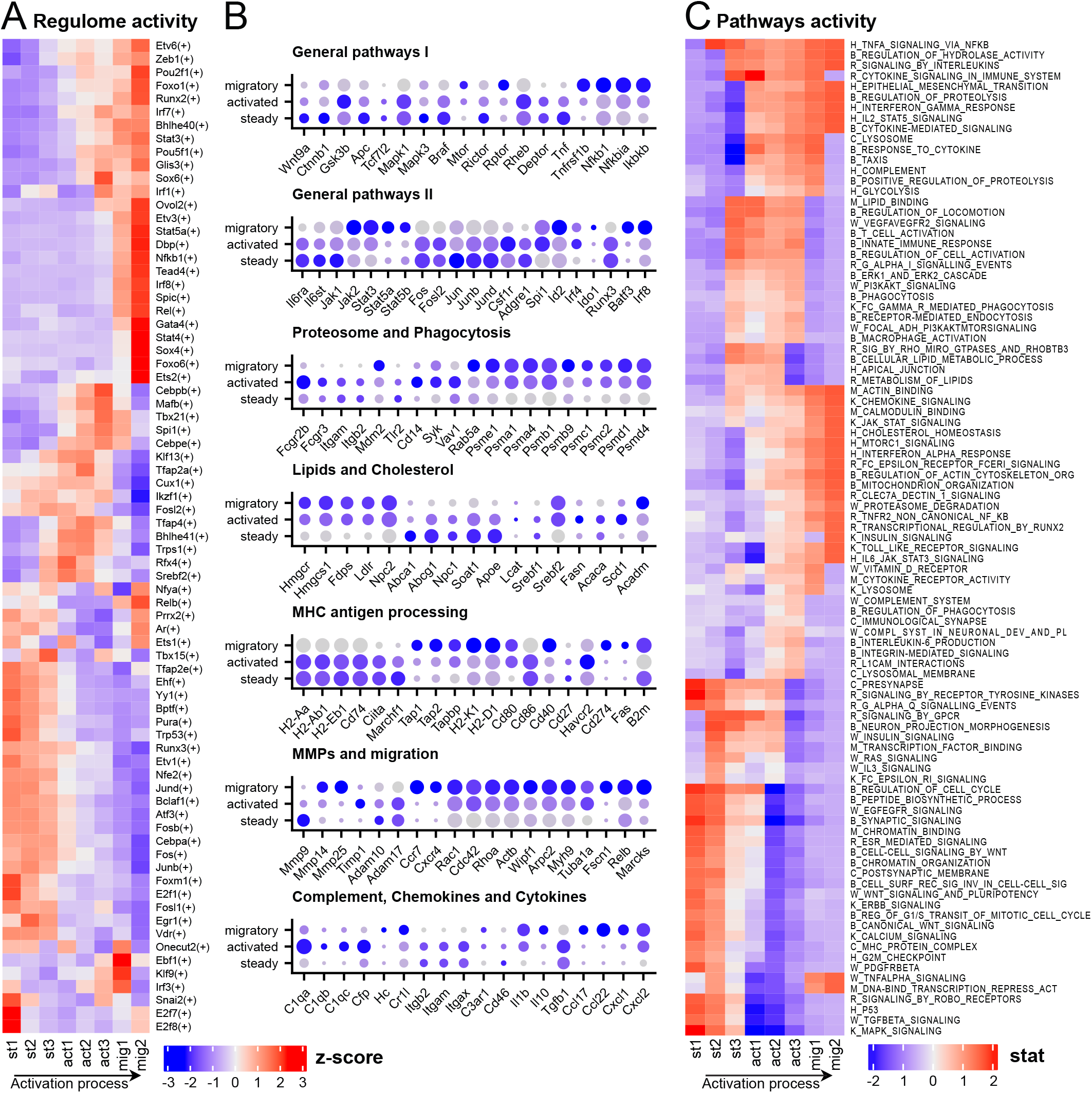
Langerhans cell (LC) activation is associated with coordinated changes in gene regulatory networks and signaling pathways. **A.** Heatmap generated using SCENIC showing the inferred activity of gene regulatory networks across LC states. Red indicates high relative activity, and blue indicates low relative activity (z-score). **B.** Dot plots showing the relative expression of selected genes across LC states. Dot size represents the proportion of cells expressing the gene, and color intensity indicates scaled expression levels. **C.** Pathway activity analysis was performed using the SCPA package. Red and blue indicate higher and lower relative pathway activity, respectively. Prefixes before pathway names denote the source database: H, Hallmarks; B, Gene ontology (GO): Biological Processes; M, GO: Molecular Functions; C, GO: Cellular Components; R, Reactome; W, WikiPathways; and K, Kyoto Encyclopedia of Genes and Genomes (KEGG).

Notably, steady-state LCs exhibit basal activity of several core signaling pathways that support homeostasis, maintain LCs within an intact epidermal niche, and sustain proliferation. These include the mitogen-activated protein kinase^[42]^, epidermal growth factor receptor^[43]^, transforming growth factor-β^[44]^, wingless-related integration site (WNT)^[45]^, and tumor protein P53^[46]^ pathways, as well as transcription factors of the Fos/Jun/activator protein 1 family^[47]^ (*Fos*, *Fosb*, *Junb*, *Jund*, *Fosl1*). Steady-state LCs exhibit transcriptional signatures related to cell-cycle progression from the pathway enrichment analysis (**Figure 3A–C**). Activated and migratory LCs exhibit mechanistic target of rapamycin (mTOR) pathway^[48]^ changes by switching from mTORC2 to mTORC1. The signaling landscape of migratory LCs is defined by the strong activation of cytoskeletal reorganization pathways, especially those involving actin dynamics controlled by Rho family GTPases^[49]^, including *Cdc42*, *Rac1*, and *RhoA*. In addition, during the transition from activation to migration, migratory LCs highly express several metalloproteases, including *Mmp14* and *Mmp25*. This suggests their role in promoting LC migration through tissues and into the lymphatic system. Activated LCs showed reduced S-phase representation compared with that of steady-state LCs, and migratory LCs were predominantly in the G1/G0 phase^[50]^. This pattern was consistent with progressive cell-cycle attenuation during activation and migration from the epidermis (**Figure 3B**). Collectively, these findings indicate a dynamic transition in the cellular state as LCs progress from tissue residency to migration.

Interestingly, state-specific changes in metabolic characteristics were observed. The negative regulator of glucose metabolism *Txnip*^[51]^ is active in steady-state LCs. In addition, transcripts of proteins responsible for metabolism and lipid transport^[52]^, such as *Apoe*, *Abcg1*, and *Abca1*, are expressed. These data suggest that steady-state LCs display transcriptional signatures consistent with fatty acid oxidation^[53]^ and moderate oxidative phosphorylation (OXPHOS)^[54]^. Moreover, activated LCs are consistent with a shift toward aerobic glycolysis, as indicated by the enrichment of the glycolytic pathway and the upregulation of key glycolytic enzymes, including *Eno1*, *Pfkfb3*, *Pfkfb4*, *Pkm*, and *Gapdh*^[55,56]^. Mitochondrial metabolism-associated transcripts increased with increased expression of *Gpd2*, *Gpx1*, *Ftl1*, and *Ehd1*, which suggests complementary oxidative support for energy and redox balance^[57–59]^. Simultaneously, lipid metabolism remained active and expanded, as highlighted by the activation of *Fabp5*, *Apol7c*, and *Cyp51* and the enrichment of the cholesterol homeostasis pathway^[60,61]^. During the transition to the migratory state, the expression of glycolysis-related genes progressively decreased, as reflected by the reduced expression of key glycolytic enzymes^[62]^, such as *Hk2*, *Gpi1*, *Pfkp*, and *Ldha*. This pattern is consistent with a relative increase in OXPHOS-associated gene expression, supported by upregulation of mitochondrial genes, including *Sdhaf1*, *Ndufa11*, *Cox5a*, and *Atp6v0e* (**Figure 3B**). Hence, our data suggest transcriptional remodeling of metabolic programs during LC activation.

Considering that LCs are key immune cells, we investigated the regulation of immunological pathways during state transitions. The expression of multiple nuclear factor-kappa B (NF-κB) pathway components increases from an early activation state and peaks at the terminal migratory state^[22]^. We found that pro- and anti-inflammatory signals coexisted during LC activation and migration. Although components of the pro-inflammatory NF-κB pathway are upregulated^[22]^, the expression of *Nfkbia*, which is an inhibitor of NF-κB, also increases^[63]^. Similarly, certain pro-inflammatory mediators, such as *Il1b*^[64]^, *Il21r*, and *Cd137* were upregulated, whereas others, including *Tnf*^[65]^, *Il1r1*, *Il13ra1*, *Il18*, and *Il1rap*, were downregulated. Simultaneously, several anti-inflammatory genes, such as *Socs2*^[66]^, *Tnfrsf11b*, *Il4i1*, *Il10*, and *Il1rn*, showed increased expression. This pattern suggests the coordinated regulation of inflammatory signaling during LC activation and migration. In addition, we observed increased expression of chemokines, such as *Ccl22*, *Ccl17,* and *Ccl5*, which are involved in recruiting T cells to lymphoid tissues^[67,68]^ (**Figure 3B**). These transcriptional patterns are consistent with a balanced immune state during LC activation and migration^[65,69,70]^.

Considering that LCs are antigen-presenting cells, we examined the expression of phagocytosis-associated genes in LCs in different states. Activated LCs express several bacterial and fungal pattern recognition receptors and key components involved in phagosome formation, including *Syk*^[71]^ and *Vav1*^[72]^. Genes encoding the activator and core subunits of the proteasome^[73]^ and immunoproteasome^[74]^, such as *Psme*, *Psma*, *Psmc*, and *Psmd*, were also expressed in the activated state. In addition, *Fcgr2b* and *Fcgr3*, which facilitate the uptake of antibody-opsonized antigens^[75]^, are upregulated in the activated state. We observed the coordinated downregulation of genes related to major histocompatibility complex (MHC) class II antigen presentation in the activated and migratory states, including *Cd74*, *Ciita*, and *Marchf1*. This downregulation may be associated with altered turnover of surface MHC-II– antigen complexes^[76]^, although its consequences for antigen presentation were not directly assessed. In contrast, components of the MHC class I pathway and cross-presentation machinery remain active in the activated and migratory states (**Figure 3B**). Overall, our results provide a detailed transcriptional view of phagocytosis- and antigen processing-associated programs across distinct LC activation states.

Moreover, genes associated with the complement system were differentially expressed during LC activation. In particular, the subunits of the C1q^[77]^ complex (*C1qa*, *C1qb*, and *C1qc*) were upregulated, which suggests the involvement of the classical complement pathway. Unexpectedly, we detected the expression of properdin (*Cfp*), which stabilizes the alternative pathway^[78]^. LCs do not express the enzymes required for the activation of the C1q complex nor the central components of the classical complement system, including C3^[79]^ and its converting enzymes. Despite the absence of full complement activation machinery, LCs express several complement receptors, including *Itgam* (*Cr3a*), *Itgax* (*Cd11c*), and *C3ar1*. This indicates their potential capacity to sense complement components. Although we observed the expression of C5 (Hc) during the late stage of migration, neither its receptors nor the enzymes required for generating anaphylatoxin C5a were detected (**Figure 3B**). These expression patterns suggested that complement-associated transcriptional programs accompany LC activation and migration, potentially in coordination with signals from other cell types.

### 2.4. Separate Clustering of the Untreated Sample Reveals Six Homeostatic LC Populations

To identify distinct homeostatic populations of LCs, we independently analyzed and clustered cells from untreated mice (**Figure S10A**). In this untreated sample, we identified six populations, with the most common population being fully quiescent LCs, followed by a prominent population characterized by a stress-responsive immunomodulatory phenotype. Differential expression analysis (**Table S2**) revealed that these immunomodulatory LCs expressed a range of heat shock proteins and chaperones (*Hsph1*, *Hspa1a*, *Hspa1b*, *Dnajb1*, *Dnajb4*, *St13*, *Bag3*, *Ubqln1*) and immunoregulatory genes (*Il18r1*, *Il1rl2*, *Tnfaip8l2*, *Tnfaip8l3*, *Slamf7*, *Card9*, *Nfatc3*, *Sirpa*). These cells were located on the opposite side of the UMAP projection relative to the pre-activated cluster; hence, they are distinct from the activated LC population. Between the homeostatic and stress-responsive immunomodulatory populations, we identified a transitional cluster marked by the expression of *Mrnip*^[80]^, which is a known activator of DNA damage response signaling. The dividing LC population expressed the full spectrum of proliferation markers (*Mki67*, *Kif11*, *Cdk1*, *Mcm3*, *H1f4*, *Stmn1*). These data were supported by CytoTRACE2 analysis (**Figure S10B**). Among the genes enriched in specific homeostatic clusters, *Pcdh7*, which is a cell adhesion protein, was differentially expressed. To assess its potential association with LC spatial organization, we analyzed epidermal whole-mounts from *Pcdh7*-deficient mice. These results indicate that *Pcdh7* contributes to the spatial organization of homeostatic LCs; however, its effect appears modest (**Figure S11A**–**C**). Overall, these results suggest that this heterogeneity reflects the diversity within homeostatic LCs and is distinct from activation-associated states.

### 2.5. Integrated Analysis Identifies Potential LC–Skin Cell Interactions

The LC-enriched dataset was integrated with the Haensel *et al.* (2020) dataset to explore potential interactions between LCs and other skin-resident cells^[81]^. The dataset includes diverse skin cell populations under unwounded and wounded conditions but contains relatively few LCs. Therefore, integration with our dataset increased LC representation within a broader skin cell context. Furthermore, ligand–receptor analysis of the integrated dataset predicted potential interactions between LCs and multiple cell populations (**Figure 4A, B**). The dataset was integrated using Seurat, and ligand–receptor interactions were identified using LIANA++ (**Table S3**), which aggregates multiple ligand–receptor interaction databases. Dermal fibroblasts and myofibroblasts were among the predicted interacting populations, including WNT-modulated fibroblast subsets^[82]^ present during tissue injury (**Figure 4C**). The key interacting receptors on LCs include *Cd44*, complement receptors, integrins, and WNT-related receptors, whereas fibroblasts predominantly express extracellular matrix components^[83]^ (**Figure 4D**). In the steady state, interactions with fibroblasts are dominated by homeostatic extracellular matrix (ECM) components, including the basement membrane. Activation induces integrin-mediated interactions between LCs and fibroblasts (e.g., *Itga9_Itgb1*) and immune-related signaling pathways (innate immunity, chemokines, and cytokines), thus reflecting ECM remodeling and immune readiness. In the migratory state, LCs shift their ECM interactions from *Cd44* to high-affinity^[84]^ integrins (*Itga9_Itgb1*, *Itga5_Itgb1*, *Itgav_Itgb1*). In myofibroblasts, the main interactions are mediated by Notch and adrenergic signaling, which increase as the LCs are activated and subsequently migrate (**Figure 4D**). Furthermore, we observed transcriptional differences between the migrating LCs and a broader DC population; however, these differences were not investigated further in this study (**Figure 4E**). Overall, these predicted ligand–receptor relationships suggest that the potential interaction landscape between LCs and multiple skin cell populations changes as LCs transition from tissue-resident to migratory states.

**Figure 4.**
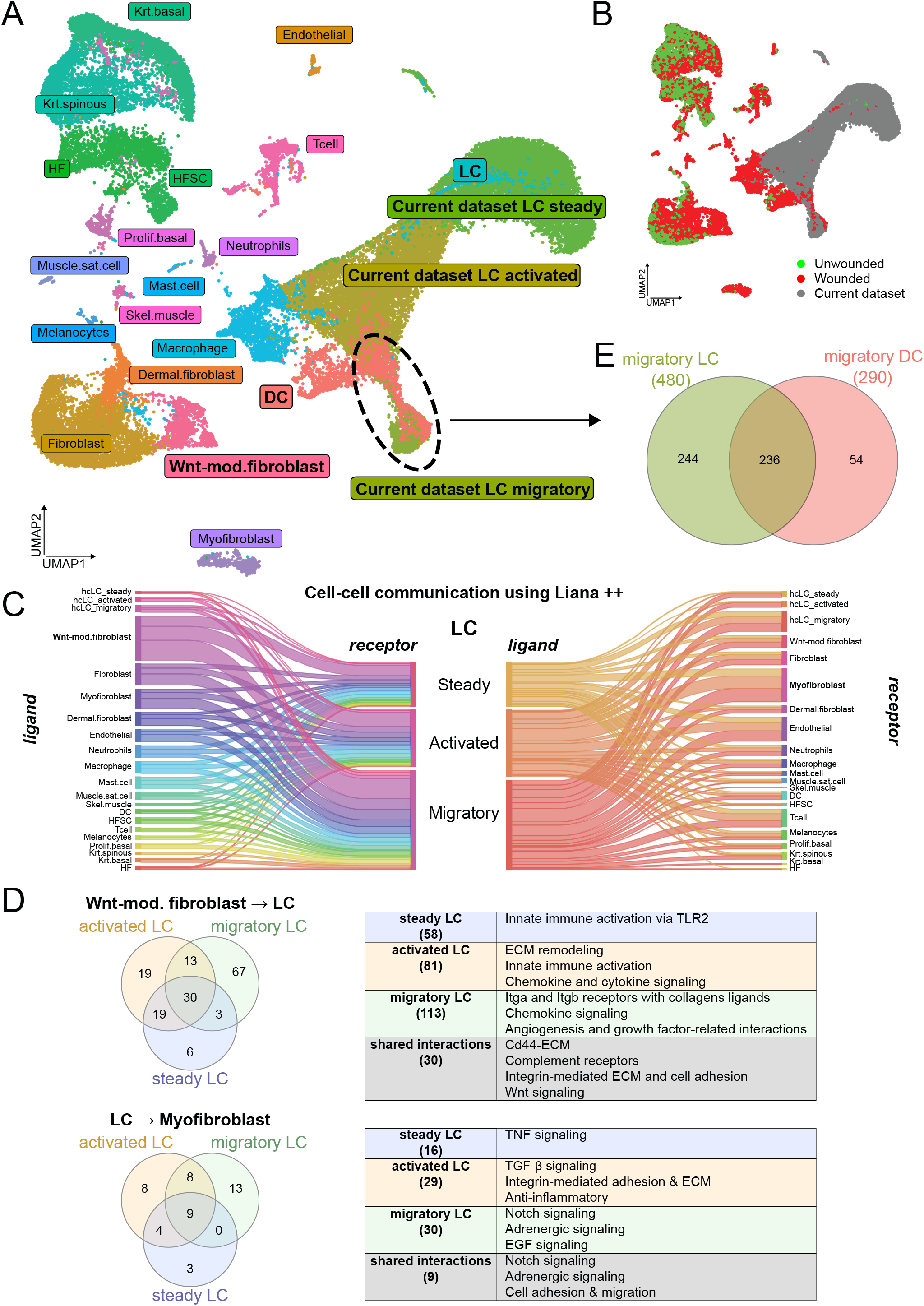
Langerhans cell (LC) state transitions are associated with dynamic changes in predicted cell–cell interactions within the skin microenvironment. **A.** Integration of the present dataset with Haensel *et al*. (2020) visualized via Uniform Manifold Approximation and Projection (UMAP), showing the relative positioning of migratory dendritic cells (DC) and migratory LCs based on transcriptional profiles. **B.** UMAP visualization of the integrated dataset, highlighting samples from Haensel *et al*. (2020): wounded skin (red), unwounded skin (green). LC data from this study are shown in grey. **C.** LIANA++ analysis predicts ligand–receptor interactions between LCs and surrounding cell populations, including wingless-related integration site (WNT)-modulated fibroblasts. **D.** Relative interaction scores between LCs and WNT-modulated fibroblasts across LC states (steady, activated, migratory). **E.** Comparative transcriptional analysis of migratory LCs and migratory DCs.

### 2.6. The Complement Cascade Contributes to LC Accumulation at Wounded Areas

To identify candidate pathways associated with the activated LC state, we examined complement-related gene expression. Transcriptomic analysis of the LCs implicated the complement system as a candidate innate immune pathway associated with LC activation (**Figure 3B**). In particular, activated LCs express C1q subunits, which are central for initiating the classical complement pathway (**Figure 5A**)^[85]^. Simultaneously, fibroblasts, especially the WNT-modulated subset^[86]^, expressed *C1s1* and *C1ra*, which are serine proteases required for the activation of C1q (**Figure 5B**). This interaction supports the initiation of the complement cascade, which promotes pathogen recognition and clearance through opsonization^[87]^. Furthermore, WNT-modulated fibroblasts expressed *C3*, which is a central component of the complement system, along with *C4b*, which is required for conversion in the complement cascade^[88]^ (**Figure 5B**, **Figure S12**). This expression pattern suggested potential complement-related interactions between LCs and fibroblasts in the local immune response.

**Figure 5.**
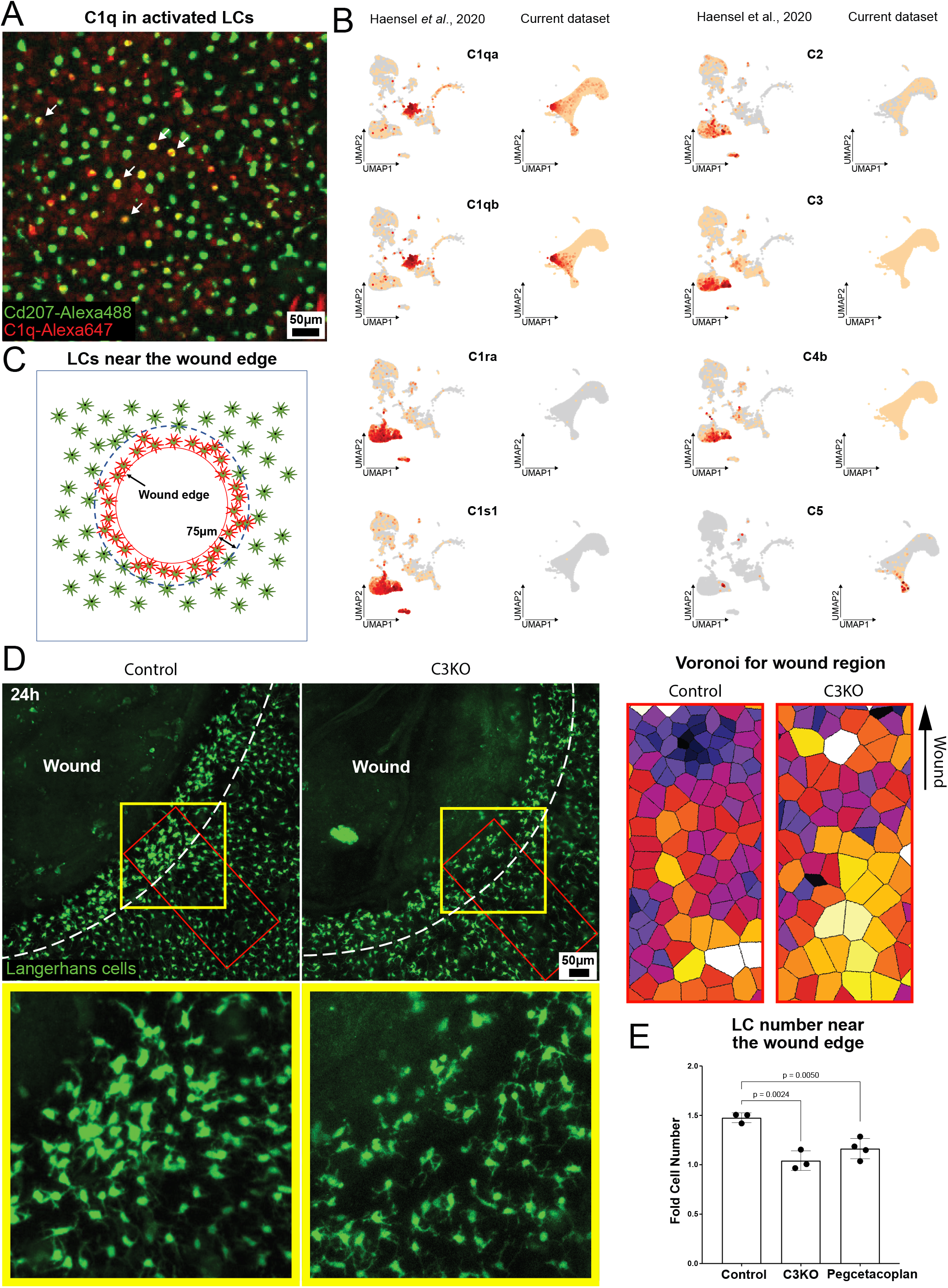
Complement signaling regulates Langerhans cell (LC) accumulation and spatial distribution following skin injury. **A.** Representative image showing *C1q* expression in LCs 48 h after croton oil/acetone treatment of ear skin; red: C1q, green: Lang-GFP. Arrows indicate C1q^+^ LCs (n = 3 mice). **B.** FeaturePlot showing expression of complement-related genes across epidermal cell types. **C.** Schematic representation of the 75-μm zone surrounding the wound edge used for quantification. **D.** Representative images and corresponding Voronoi diagrams of LCs near the wound edge in control and *C3* knockout (C3KO) mice (n = 3 mice). The yellow box indicates the area enlarged below, the red box indicates the region used for the Voronoi analysis shown on the right, and the dotted line marks the 75-μm zone from the wound edge used for LC quantification. **E.** Quantification of LC enrichment within the 75-μm circumferential zone around the wound in control, C3KO, and pegcetacoplan-treated mice. Fold change in LC number at 24 h post-injury was calculated relative to values immediately after wound creation in the same mouse. Statistical analysis was performed using a paired t-test (n = 3, 4 mice). Bar plot statistics were calculated using a paired t-test.

One *C3* derivative is anaphylatoxin C3a, which attracts antigen-presenting cells to sites of inflammation. Although LCs express an anaphylatoxin receptor, this observation has not been described in the context of LC function^[89]^. Therefore, to investigate the functional role of *C3* in complement activation and LC localization, *C3*-deficient (C3KO) mice combined with the *Lang-GFP* allele were generated to visualize LCs using intravital imaging. For spatial quantification, 1-mm circular ear wounds were generated. Unlike the broadly applied injury models used above, this approach provides a defined wound edge for reproducible measurement of LC accumulation (**Figure 5C**). In control mice, LCs lost their regular distribution and were enriched near the wound edge 24 h after injury. In contrast, C3KO mice exhibited reduced LC enrichment at the wound edges (**Figure 5D**). Pharmacological inhibition of C3 with pegcetacoplan, a pegylated C3/C3b inhibitor^[90]^, recapitulated the phenotype observed in C3KO mice, with a cell density close to the baseline levels near the wound (**Figure 5E**). These results indicate that C3 contributes to LC accumulation and localization at the injury site. Collectively, our results suggest that LC behavior following injury is associated with signals from the surrounding tissue microenvironment, including complement-related cues (**Figure S13**).

## 3. Discussion

LCs are tissue-resident immune cells in the epidermis that play an important role as a first-line immune barrier to the external environment. Upon activation by various external stimuli, LCs migrate to the lymphatic system to initiate adaptive immune responses. However, the transcriptomic dynamics and functional consequences of LC activation remain poorly understood. In this study, scRNA-seq of murine epidermal LCs enabled the investigation of LC transcriptional diversity and function-associated transcriptional signatures. Although mRNA levels do not always directly mirror protein abundance^[91,92]^, scRNA-seq remains the most informative and scalable approach for resolving cellular states *in vivo* and provides a high-resolution framework for interpreting function-associated transcriptional heterogeneity^[93,94]^. LCs transition from a steady state to a migratory state upon activation and previous studies have treated this process as a binary switch. Furthermore, heterogeneity exists within LC populations; however, the intermediate stages of activation and their functional characteristics remain poorly defined. In this study, we extended these findings by identifying a distinct intermediate activation state and reconstructing a trajectory-resolved program for LC activation *in vivo*. This framework provides a more detailed view of the LC state transitions and their associated transcriptional changes during tissue injury. In this study, we obtained a high-quality single-cell dataset of over 22,000 LCs, which encompassed the entire activation trajectory. Our data captured the dynamic progression of LCs from a homeostatic steady state through a previously unrecognized intermediate activated state to a fully activated migratory state.

The scRNA-seq analysis combined with RNA velocity analysis revealed that LCs exhibit similar activation trajectories in response to both physical (stamp punch) and chemical (croton/acetone) stimuli, thereby indicating overlapping LC transcriptional responses in these damage-driven models. Mechanistically distinct stimuli, such as UV irradiation or contact sensitizers (e.g., DNFB), which elicit distinct LC responses, have not been examined and may engage in different transcriptional programs. Using intravital imaging, we confirmed that the LCs underwent similar spatial redistribution at comparable time points after both treatments. Considering that the 46-h time point was selected based on the spatial phenotype, our single-time-point scRNA-seq analysis does not establish when the activated state was most abundant transcriptionally. Earlier or later sampling may alter the relative representations of steady, activated, and migratory LC states. Nevertheless, the selected time point enabled effective capture and characterization of an intermediate activated LC state between the steady and migratory states. Moreover, we assembled a detailed map of the molecular features of the activated state separately from those of the migratory and steady states. Specifically, activated LCs show enrichment of glycolysis-associated transcripts, upregulation of complement system components, and increased expression of phagocytosis-related genes. Activated LCs also upregulate *Cd14*, which is a marker that is not classically associated with LCs. This occurs in cells that retain their canonical LC identity and coincides with the induction of phagocytosis-related genes. The significance of *Cd14* expression in this context remains unclear. However, it may reflect an activation-associated transcriptional state, which warrants further investigation. These transcriptional changes may reflect the distinct cellular programs associated with LC activation and subsequent migration. In addition, specific markers, such as *Cd137*, *Ly6a*, and *Cd14,* reliably distinguish activated-state LCs, thus providing an unprecedented basis for convenient activated LC sorting in future studies. The use of these markers was consistent with the activated LC transcriptomic state; however, additional molecular validation of the sorted populations would further strengthen this link.

Our analysis revealed that LCs transition through three distinct states: steady, activated, and migratory. Furthermore, these states have distinct transcriptional signatures associated with metabolic and immunological processes. In particular, steady-state LCs are enriched in homeostasis- and fatty acid oxidation-associated programs, whereas activated LCs show increased expression of genes associated with glycolysis, phagocytosis, and antigen processing. Correspondingly, migratory LCs are enriched in genes associated with oxidative metabolism, cytoskeletal remodeling, and tissue egress. These coordinated transcriptional changes distinguish between the steady, activated, and migratory LC states. However, further *in vivo* studies are required to determine the role of these state-associated programs in LC function.

Notably, our findings highlight the complement system as a key regulator of LC behavior during skin inflammation. Specifically, the spatially coordinated expression of complement components, where fibroblasts serve as local sources of C3 and its activating enzymes and LCs express complement receptors, suggests potential complement-related signaling between fibroblasts and LCs. Furthermore, impaired LC recruitment observed in C3-deficient mice supports a role for C3 in LC positioning. Importantly, pharmacological inhibition of C3 using pegcetacoplan recapitulated the phenotype observed in C3 knockout mice. This further supports a role for C3-dependent signaling in regulating LC positioning near the wound. Collectively, these data support the association between complement signaling and LC positioning following tissue injury.

Furthermore, we observed expression of *C5* in LCs transitioning toward the migratory state (**Figure S12**). However, the analysis of major skin cell populations showed that *C5* expression was not restricted to LCs and was detectable in other cell populations, including fibroblast subsets. Therefore, the cellular source and functional contribution of *C5* during skin inflammation remain to be determined. Considering the established association of C5 cleavage products with inflammatory skin diseases, such as psoriasis and atopic dermatitis^[95–100]^, these observations suggest that C5-associated complement signaling may contribute to the local inflammatory environment during skin injury.

In conclusion, we defined a distinct activation trajectory of epidermal LCs and identified a transcriptionally distinct intermediate state characterized by the enrichment of phagocytosis- and complement-associated gene expression. Our findings indicate that LCs exhibit distinct transcriptional programs across activation states and provide a framework for understanding LC state transitions during skin injury and immune responses.

## 4. Methods

### 4.1. Mice

CD-1 mice were obtained from Charles River Laboratories. Lang-EGFP (JAX 016939) and C3KO (JAX 029661) mice were obtained from the Jackson Laboratory. K14-H2BmCherry mice were obtained from V. Greco (Yale School of Medicine). *Pcdh7*-knockout mice were obtained from Dr. Yongwon Choi (University of Pennsylvania). To simultaneously visualize LCs and epithelial cells, Lang-EGFP; K14H2B-mCherry mice were generated. To knock out C3 and visualize LCs, the Lang-EGFP; K14H2B-mCherry, and C3 KO^−/−^ mice were generated. Siblings were used as controls (Lang-EGFP; K14H2B-mCherry; C3 KO^+/−^). Mice were housed in ventilated racks under controlled conditions, including an ambient temperature of 22 °C, relative humidity of 50 ± 10%, and a 12-h light/dark cycle (lights on from 07:00 to 19:00). Male and female mice from experimental and control groups were randomly selected for live imaging and LC isolation. All animals used in this study were between five and eight weeks of age. No blinding was conducted. All animal procedures were approved by the Institutional Animal Care and Use Committee (IACUC) at Michigan State University under

PROTO202600073 and were conducted in accordance with the institutional and national guidelines for animal welfare.

### 4.2. *In Vivo* LC Activation

The mice were anesthetized in an isoflurane chamber, and anesthesia was maintained throughout the experiment using vaporized isoflurane delivered via a nose cone, as previously described^[101]^. Fur was removed from the ears and head using Nair cream (Naircare; 022600003014) applied for 2 min using a Q-tip, followed by tap water cleaning and a second 1-min application. To prevent physical damage to the skin, electric razors were not used. The croton oil solution was freshly prepared before each treatment by mixing croton oil (Sigma; C6719), acetone (Sigma; 179124), and olive oil (Sigma; O1514) in a 4:5:1 ratio. Mouse ears were carefully affixed and flattened onto plastic cubes (15 × 15 mm) using petroleum jelly (Vaseline). Additional petroleum jelly was applied to the base of the ear (proximal to the head) to prevent solution leakage. A total of 35 μL of the croton oil mixture was applied to the external surface of each ear and evenly distributed using a plastic pipette tip. Both ears were simultaneously treated. Following the application, the mice were left undisturbed for 10 min; the mice were anesthetized using an isoflurane mask during the procedure. Thereafter, the ears were detached from the plastic cube, and the mice were returned to the isoflurane chamber for an additional 10 min. This procedure was repeated 24 h later. Croton oil-activated LCs were isolated 48 h after the initial treatment. For physical treatment using microneedle puncture, a microneedle array consisting of 38 needles, each 0.2 mm in length (Skinmedix; micro needle dermal stamp 0.2 mm; 765573893595), was used. Fur removal and ear fixation were performed as described above. A total of 750 neat microneedle punctures were applied without pressing to each ear and evenly distributed across the ear surface; the mice were anesthetized using an isoflurane mask during the procedure. The procedure was repeated 24 h later with a reduced number of punctures (150 punctures per ear). LCs were isolated from the stamp-punched skin 48 h after the first treatment.

### 4.3. Epidermal Single-Cell Suspension Preparation and LC Sorting

Mice were treated as described above, whereas control mice received only Nair treatment at the same time as the experimental animals. The procedure was performed 48 h after the initial treatment in both croton oil and microneedle punch. The mice were euthanized following standard protocols after being anesthetized with isoflurane. Subsequently, the ears were excised, and the skin layer was carefully separated from the underlying cartilage. The isolated skin was placed epidermis-side up in a trypsin solution (0.3% trypsin (Thermofisher; 15090046) in 150 mM NaCl, 0.5 mM KCl, and 0.5 mM glucose) and incubated for 1 h 45 min at 37 °C. After incubation, the epidermis was gently separated from the dermis using fine forceps. The epidermal sheet was transferred to a scratched-wall plastic Petri dish (Fisher; FB0875711YZ) containing Hanks’ Balanced Salt Solution supplemented with 10% fetal bovine serum, trypsin inhibitor (Sigma; T6522), and DNaseI (Sigma; D4513). Thereafter, the sheet was pre-warmed to 37 °C. The epidermis was mechanically dissociated into a single-cell suspension by pressing it against a scratched surface using forceps to produce a cloudy cell suspension. This suspension was incubated for 15 min at 37 °C and then filtered through a 70-µm cell strainer. Dead cells were removed via magnetic separation (Miltenyibiotec; 130-090-101), followed by the magnetic enrichment of LCs (Miltenyibiotec; 130-095-408). Cell viability was assessed via trypan blue staining or flow cytometry (Protocol S1).

### 4.4. Single-Cell Library Preparation and Sequencing

For scRNA-seq, libraries were prepared according to the manufacturer’s manual using the Chromium Next GEM Single-Cell 3’ Kit v3.1 (10x Genomics, USA) and sequenced using two lanes (2.5GB) NovaSeq X 10 B 100 SR (Illumina, USA).

### 4.5. Single-Cell Data Analysis

#### 4.5.1. Preprocessing

Raw scRNA-seq data were processed using Cell Ranger (v8.0, 10x Genomics) with default parameters and aligned to the GRCm39 reference genome. This was customized to include additional coding sequences for mCherry (AY678264.1) and eGFP (OQ870305.1).

#### 4.5.2. Quality Control and Clustering

Quality control, normalization, clustering, and visualization were performed using Seurat^[102]^ (v5.2) in R (4.4.0) and Scanpy^[103]^ (v1.10) with Python 3.10 for RNA velocity and cell–cell communication. In our quality control process, cells were filtered based on mitochondrial content and gene/UMI counts using the following Seurat parameters: nFeature_RNA > 1500 & nFeature_RNA < 7000 & nCount_RNA > 1500 & nCount_RNA < 40000 & percentage mt < 3.5. Data were normalized using NormalizeData, clustered using the Louvain algorithm, and visualized using UMAP. Considering the homogeneity of LC data, UMAP was performed using the top five principal components. To avoid integration-related artifacts in our relatively homogeneous and high-quality dataset, we initially analyzed the LC data without applying advanced integration algorithms. The resulting clustering pattern, including the division into three distinct clusters of LCs, was subsequently validated via integration with the dataset from Haensel *et al*. (2020). For various helper functions, scCustomize^[104]^ was used.

#### 4.5.3. Data Integration

To integrate the Haensel *et al*. (2020) data, Seurat’s standard integration workflow was used. Anchors were identified across datasets using FindIntegrationAnchors, followed by IntegrateData to obtain a corrected expression matrix that was used for downstream clustering and visualization. The two datasets were generated in separate experimental settings; hence, the predicted ligand–receptor relationships were interpreted as exploratory and hypothesis-generating rather than evidence of direct cell–cell communication.

#### 4.5.4. Pathway Analysis

SCPA^[105]^ (v1.6.1) was used to compute single-cell pathway activity scores using curated gene sets (MSigDB^[106,107]^, Reactome^[108]^, and Wikipathways^[109]^).

#### 4.5.5. Gene Regulatory Networks

SCENIC^[110]^ (v0.12.1) was used to infer transcription factor activity and regulons using GENIE3 (v1.28), RcisTarget (v1.26), and AUCell (v1.28).

#### 4.5.6. RNA Velocity

Velocyto^[111]^ (v0.17.15) and scVelo^[112]^ (v0.2.5) were used for RNA velocity analysis, while employing the dynamical model and projecting velocities onto UMAP.

#### 4.5.7. Trajectory Analysis

Trajectory inference was performed using Slingshot^[113]^ (v2.14.0) with UMAP embeddings and Seurat cluster annotations as input. The steady-state LC cluster was defined as the root and the pseudotime values were computed without predefining the terminal states. Genes dynamically regulated along the trajectory were identified using tradeSeq^[114]^ (v1.20.0). This system fits generalized additive models to detect significant expression changes across pseudotime (adjusted P < 0.05).

#### 4.5.8. Cellular Potency Analysis

The cellular potency of the control sample was predicted using CytoTRACE2^[115]^, which infers differentiation potential from single-cell transcriptomic profiles based on gene expression diversity and associated transcriptional features.

#### 4.5.9. Cell Cycle

Cell-cycle phase distribution was assessed using the scCustomize::Add_Cell_QC_Metrics function, which estimates the proportion of cells in the G1, S, and G2/M phases based on canonical gene signatures. The resulting annotations were integrated into the Seurat object and visualized using dittoSeq::dittoBarPlot to compare the cell-cycle composition across LC states. This analysis was performed to evaluate whether the observed transcriptional differences were associated with cell-cycle variations.

#### 4.5.10. Cell–Cell Communication

LIANA++^[116]^ (v0.1.12) was used to infer ligand–receptor interactions across cell types using consensus rankings from multiple methods.

### 4.6. Intravital Imaging

Mice were anesthetized using an isoflurane induction chamber, and anesthesia was maintained throughout imaging via a nose cone delivering vaporized isoflurane, as previously described. Imaging was performed using a Leica TCS SP8 X Two-Photon microscope equipped with a Spectra Physics Insight MP Laser and controlled using Leica LAS X software. Z-stack images were acquired using a 25× water-immersion objective (NA 1.0; Leica), with a field of view of 0.59 × 0.59 mm. A wavelength of 940 nm was used to excite the LCs in Lang-EGFP mice. Z-stacks were acquired in 2-µm steps to cover depths up to 80 µm. For large-area imaging, 9–12 tiled optical sections were acquired using a motorized stage with a custom Fiji^[117]^ (v 1.53) stitching script available on GitHub (see the Data Availability section). Automatic cell segmentation was performed using Cellpose^[118]^, which was trained on multiple LC images.

### 4.7. Voronoi Diagram

The spatial distribution of the cells was analyzed using Voronoi tessellation in Fiji with the Voronoi plugin. Binary masks of the cell centroids were generated, and Voronoi diagrams were computed. The area of each Voronoi cell was measured and used to assess the spatial dispersion and local cell density.

### 4.8. Flow Cytometry

For flow cytometric analysis of LC markers, staining was performed prior to the 70-µm filtration step. Draining mandibular lymph nodes (LN) were isolated as previously described^[119]^. LN cell suspensions were prepared using the same procedure as for LCs, except for the trypsin digestion step. LNs were mechanically dissociated by pressing the tissue against a scratched surface using forceps to produce a cloudy cell suspension. After preparing the cell suspension from the epidermis or LNs, the desired number of cells was aliquoted into a 96-well polypropylene v-bottom plate (Corning; 3344). Dead cells were stained using Live/Dead Blue dye (Thermo Fisher Scientific; L34961; 1:800) for 15 min at room temperature, followed by two washes with the staining buffer. To block nonspecific binding, Fc receptors were blocked using Fc block reagent (BDbiosciences; 553142) for 15 min. Next, the cells were incubated with non-conjugated, fluorochrome-, or biotin-conjugated primary antibodies (1:200 dilution) for 30 min in the dark (BioLegend: 108103, 123305, 104703, 146511, 100212, 100425, 104545, 103228). After two additional washes with the staining buffer, the cells were incubated with streptavidin-conjugated fluorochrome (1:800 dilution) or the appropriate fluorochrome-conjugated secondary antibodies (BioLegend: 405237). Following staining, the cells were washed several times with the staining buffer. The cells were then fixed with 4% paraformaldehyde in phosphate-buffered saline (PBS) for 15 min, washed three times with flow cytometry buffer, and analyzed using flow cytometry. Flow cytometry was performed using the Cytek Aurora spectral flow cytometer. Cells were acquired using the SpectroFlo software. Compensation and unmixing were performed automatically based on single-stain controls, using an autofluorescence exclusion function. Data were exported to FCS 3.1. Flow cytometry data were analyzed using FCS Express (v 7.24, De Novo Software). The initial gating was performed to exclude debris, doublets, and dead cells. Subsequent gating strategies were applied based on marker expression to identify cell populations of interest. Data visualization included biaxial plots and population statistics.

### 4.9. OVA Treatment

To deliver ovalbumin (OVA) to LCs, OVA (InvivoGen; vac-stova, endotoxin-free) was diluted in PBS to a concentration of 10 mg/mL. Mice were anesthetized and immobilized, and the fur was removed from the ear and head, which was then flattened and secured onto a plastic support cube, as described above. A total of 70 µL of 100% absolute ethanol (Sigma; 459836) was applied to each ear surface. Immediately afterward, 30 µL of the OVA–PBS solution was applied and evenly distributed across each ear. The ear with the ethanol–OVA mix was dried for 10 min using a stream of argon gas to fix the protein onto the skin surface. After drying, a stable OVA film formed on the skin surface, after which a mixture of croton oil, acetone, and olive oil (prepared as described above) was applied to the ears and left in place for 10 min. Mice were then transferred to an isoflurane chamber for 10 min. For topical treatment, a 10 × 20 mm bath was formed on the lateral side of the mouse using adhesive tape. The edges of the tape were sealed with petroleum jelly to prevent leakage and ensure localized application. OVA and the croton oil mixture were applied in the same manner as for the ears.

### 4.10. Pegcetacoplan Treatment

Pegcetacoplan acetate (MedChemExpress; HY-P3252A) was prepared according to the manufacturer’s instructions. The compound was sequentially dissolved in 10% dimethyl sulfoxide (DMSO; Sigma; D2650), 40% PEG300 (MCE; HY-Y0873), and 5% Tween-80 (Sigma; P1754), followed by 45% PBS, to obtain a final working concentration of 10 mg/mL. A total volume of 15 µL (150 µg) of the working solution was applied topically to each freshly generated wound. The solution was allowed to dry under a gentle stream of argon. Control mice received a vehicle solution (10% DMSO, 40% PEG300, 5% Tween-80, 45% PBS) without pegcetacoplan, and the wounds were similarly dried with argon.

### 4.11. Whole-Mount Staining

Permeabilized whole-mount staining of full-thickness skin was performed in 24-well plates, following a previously described protocol^[120]^ using an anti-C1q antibody (Abcam: ab182451) and Alexa633 secondary (Invitrogen: A-21070).

### 4.12. Enzyme-Linked Immunosorbent Assay

Mouse serum was isolated using the gel centrifugation method and frozen for simultaneous analysis using a standard mouse anti-OVA IgG enzyme-linked immunosorbent assay following the manufacturer’s protocol (Chondrex; 3011).

### 4.13. Skin Wound

To create a round wound, a punch biopsy tool with a circular blade (diameter = 1 mm) was used (Miltex Instruments 33-31AA). The mice were anesthetized with isoflurane as previously described. A circular wound was made with a single, light motion, while carefully avoiding penetration into the dermis. The epidermal layer within the wound was gently removed using tweezers.

### 4.14. Statistical Analysis

Data are presented as mean ± standard deviation, with biological sample sizes (n) indicated in the corresponding figure legends. Unless otherwise specified, n represents individual mice, and the pooled samples are explicitly indicated. No data points were excluded as statistical outliers. Quantitative comparisons were performed using unpaired two-sided Student’s t-tests, where indicated, and P < 0.05 was considered statistically significant. For flow cytometry analyses, two-tailed Welch’s t-tests or paired t-tests were used as indicated in the figure legends, with Holm correction for multiple comparisons. Statistical analyses and graphical presentations were performed using GraphPad Prism v11.

For scRNA-seq, epidermal LCs pooled from five mice were used for each experimental condition. Cells were filtered using predefined quality-control thresholds (1,500 < nFeature_RNA < 7,000; 1,500 < nCount_RNA < 40,000; mitochondrial transcripts < 3.5%) and normalized using Seurat. Differential expression was assessed using the Wilcoxon rank-sum test with Benjamini–Hochberg correction and an adjusted P < 0.05 was considered significant. Pseudotime-associated genes were analyzed using tradeSeq with Benjamini– Hochberg adjusted P < 0.05. Single-cell analyses were performed in R v4.4.0 and Python v3.10 using the software packages specified above.

## Author Contributions

A.K.: Conceptualization, Investigation, Writing – original draft, Writing – review & editing; A.D.S.-S., S.M., and H.K.: Investigation, Mouse breeding; S.W.: Investigation; S.P.: Conceptualization, Writing – review & editing, Supervision, Funding acquisition.

## Supporting information

Supplemental Protocol 1

## Acknowledgements

We thank MSU IQ and MSU Flow Cytometry Core for their assistance with flow cytometry training and access to the Cytek Aurora FLOW cytometer. We also thank the MSU Genomics Core for library preparation. We thank Sergey S. Fomenko (Genoanalytica, Moscow) for proposing the use of pegcetacoplan as a functional control for the *C3* knockout. We thank Dr. Kim Hyunsoo and Dr. Yongwon Choi (University of Pennsylvania) for providing *Pcdh7-*knockout mice. We thank Dr. Luisa Valencia and Dr. Joao Pedra (University of Maryland) for their expert assistance in designing the FACS antibody panel for T and B cells. We also thank the University of California, Berkeley and QB3 Genomics Core for library sequencing. We thank Nicholas Basista and Audrey Bench for their technical contributions to the mouse genotyping. Illustrations were created using R studio (RStudio 2024.09.1+394), Inkscape 1.3.2, Bio-Render, GraphPad Prism (v9), and FCS Express (v7). This work was supported by the National Institutes of Health (NIH) under award numbers R01AR083086 and R01AI134696, by the MSU Discretionary Funding Initiative, and by the Department of Medicine Seed Funding.

## Data Availability Statement

All sequencing data generated in this study have been deposited in the Gene Expression Omnibus (GEO) under the accession number GSE296148. Any additional information required to reanalyze the data reported in this study is available from the lead contact upon request.

## References

1. D. H. Kaplan, “Ontogeny and Function of Murine Epidermal Langerhans Cells,” Nature Immunology 18, no. 10 (2017): 1068–1075, 10.1038/ni.3815.

2. J. Deckers, H. Hammad, and E. Hoste, “Langerhans Cells: Sensing the Environment in Health and Disease,” Frontiers in Immunology 9 (2018): 93, 10.3389/fimmu.2018.00093.

3. Z.-C. Wang, Y.-Y. Hu, X. Z. Shen, and W.-Q. Tan, “Absence of Langerhans Cells Resulted in Over-Influx of Neutrophils and Increased Bacterial Burden in Skin Wounds,” Cell Death & Disease 15, no. 10 (2024): 760, 10.1038/s41419-024-07143-1.

4. R. Wasko, K. Bridges, R. Pannone, et al., “Langerhans Cells Are Essential Components of the Angiogenic Niche during Murine Skin Repair,” Developmental Cell 57, no. 24 (2022): 2699–2713.e5, 10.1016/j.devcel.2022.11.012.

5. A. Rajesh, G. Stuart, N. Real, et al., “Depletion of Langerin^+^ Cells Enhances Cutaneous Wound Healing,” Immunology 160, no. 4 (2020): 366–381, 10.1111/imm.13202.

6. C. L. Bennett, F. Fallah-Arani, T. Conlan, et al., “Langerhans Cells Regulate Cutaneous Injury by Licensing CD8 Effector Cells Recruited to the Skin,” Blood 117, no. 26 (2011): 7063–7069, 10.1182/blood-2011-01-329185.

7. N. Nikolić, P. Kienzl, P. Tajpara, M. Vierhapper, J. Matiasek, and A. Elbe-Bürger, “The Antiseptic Octenidine Inhibits Langerhans Cell Activation and Modulates Cytokine Expression upon Superficial Wounding with Tape Stripping,” Journal of Immunology Research 2019 (2019): 1–11, 10.1155/2019/5143635.

8. E. Peterman, E. J. A. Quitevis, C. E. A. Goo, and J. P. Rasmussen, “Rho-Associated Kinase Regulates Langerhans Cell Morphology and Responsiveness to Tissue Damage,” Cell Reports 43, no. 5 (2024): 114208, 10.1016/j.celrep.2024.114208.

9. S. Holzmann, C. H. Tripp, M. Schmuth, et al., “A Model System Using Tape Stripping for Characterization of Langerhans Cell-Precursors In Vivo,” Journal of Investigative Dermatology 122, no. 5 (2004): 1165–1174, 10.1111/j.0022-202X.2004.22520.x.

10. E. E. Vine, P. J. Austin, T. R. O’Neil, et al., “Epithelial Dendritic Cells vs. Langerhans Cells: Implications for Mucosal Vaccines,” Cell Reports 43, no. 4 (2024): 113977, 10.1016/j.celrep.2024.113977.

11. L. Zhou, A. Jiang, J. Veenstra, D. M. Ozog, and Q.-S. Mi, “The Roles of Skin Langerhans Cells in Immune Tolerance and Cancer Immunity,” Vaccines 10, no. 9 (2022): 1380, 10.3390/vaccines10091380.

12. D. Y. Kitashima, T. Kobayashi, T. Woodring, et al., “Langerhans Cells Prevent Autoimmunity via Expansion of Keratinocyte Antigen-Specific Regulatory T Cells,” eBioMedicine 27 (2018): 293–303, 10.1016/j.ebiom.2017.12.022.

13. J. Davies, S. Sirvent, A. F. Vallejo, et al., “Transcriptional Programming of Immunoregulatory Responses in Human Langerhans Cells,” Frontiers in Immunology 13 (2022), 10.3389/fimmu.2022.892254.

14. G. Reynolds, P. Vegh, J. Fletcher, et al., “Developmental Cell Programs Are Co-Opted in Inflammatory Skin Disease,” Science (New York, N.Y.) 371, no. 6527 (2021): eaba6500, 10.1126/science.aba6500.

15. S. M. Tamminga, M. M. Van Der Wal, E. S. Saager, et al., “Single-Cell Sequencing of Human Langerhans Cells Identifies Altered Gene Expression Profiles in Patients with Atopic Dermatitis,” ImmunoHorizons 9, no. 2 (2025): vlae009, 10.1093/immhor/vlae009.

16. K. M. Bertram, R. A. Botting, H. Baharlou, et al., “Identification of HIV Transmitting CD11c+ Human Epidermal Dendritic Cells,” Nature Communications 10, no. 1 (2019): 2759, 10.1038/s41467-019-10697-w.

17. K. M. Bertram, T. R. O’Neil, E. E. Vine, H. Baharlou, A. L. Cunningham, and A. N. Harman, “Defining the Landscape of Human Epidermal Mononuclear Phagocytes,” Immunity 56, no. 3 (2023): 459–460, 10.1016/j.immuni.2023.02.001.

18. V. Pena-Cruz, L. M. Agosto, H. Akiyama, et al., “HIV-1 Replicates and Persists in Vaginal Epithelial Dendritic Cells,” The Journal of Clinical Investigation 128, no. 8 (2018): 3439–3444, 10.1172/JCI98943.

19. A. Kazmi, R. Gill, P. Restrepo, and A. L. Ji, “The Spatial and Single-Cell Landscape of Skin: Charting the Multiscale Regulation of Skin Immune Function,” Seminars in Immunology 78 (2025): 101958, 10.1016/j.smim.2025.101958.

20. H. Chen, F. Ye, and G. Guo, “Revolutionizing Immunology with Single-Cell RNA Sequencing,” Cellular and Molecular Immunology 16, no. 3 (2019): 242–249, 10.1038/s41423-019-0214-4.

21. S. Park, C. Matte-Martone, D. G. Gonzalez, et al., “Skin-Resident Immune Cells Actively Coordinate Their Distribution with Epidermal Cells during Homeostasis,” Nature Cell Biology 23, no. 5 (2021): 476–484, 10.1038/s41556-021-00670-5.

22. K. Clayton, A. F. Vallejo, J. Davies, S. Sirvent, and M. E. Polak, “Langerhans Cells— Programmed by the Epidermis,” Frontiers in Immunology 8 (2017): 1676, 10.3389/fimmu.2017.01676.

23. B. Z. Igyártó, and D. H. Kaplan, “Antigen Presentation by Langerhans Cells,” Current opinion in immunology 25, no. 1 (2013): 115–119, 10.1016/j.coi.2012.11.007.

24. A. Kissenpfennig, S. Henri, B. Dubois, et al., “Dynamics and Function of Langerhans Cells in Vivo: Dermal Dendritic Cells Colonize Lymph Node Areas Distinct from Slower Migrating Langerhans Cells,” Immunity 22, no. 5 (2005): 643–654, 10.1016/j.immuni.2005.04.004.

25. G. Mao, D. Douglas, M. Prajapati, et al., “Investigation of Inflammatory Mechanisms Induced by Croton Oil in Mouse Ear,” Current Research in Toxicology 7 (2024): 100184, 10.1016/j.crtox.2024.100184.

26. S. C. Zheng, G. Stein-O’Brien, L. Boukas, L. A. Goff, and K. D. Hansen, “Pumping the Brakes on RNA Velocity by Understanding and Interpreting RNA Velocity Estimates,” Genome Biology 24, no. 1 (2023): 246, 10.1186/s13059-023-03065-x.

27. J. Valladeau, V. Clair-Moninot, C. Dezutter-Dambuyant, et al., “Identification of Mouse Langerin/CD207 in Langerhans Cells and Some Dendritic Cells of Lymphoid Tissues,” Journal of Immunology (Baltimore, Md.: 1950) 168, no. 2 (2002): 782–792, 10.4049/jimmunol.168.2.782.

28. L. Li, L. Yang, and D. Jiang, “Research Progress of CD80 in the Development of Immunotherapy Drugs,” Frontiers in Immunology 15 (2024): 1496992, 10.3389/fimmu.2024.1496992.

29. H. Yokozeki, K. Takayama, O. Ohki, et al., “Comparative Analysis of CD80 and CD86 on Human Langerhans Cells: Expression and Function,” Archives of Dermatological Research 290, no. 10 (1998): 547–552, 10.1007/s004030050350.

30. K. Kabashima, N. Shiraishi, K. Sugita, et al., “CXCL12-CXCR4 Engagement Is Required for Migration of Cutaneous Dendritic Cells,” The American Journal of Pathology 171, no. 4 (2007): 1249–1257, 10.2353/ajpath.2007.070225.

31. E. J. Villablanca, and J. R. Mora, “A Two-Step Model for Langerhans Cell Migration to Skin-Draining LN,” European journal of immunology 38, no. 11 (2008): 2975–2980, 10.1002/eji.200838919.

32. X. Liu, R. Zhu, Y. Luo, et al., “Distinct Human Langerhans Cell Subsets Orchestrate Reciprocal Functions and Require Different Developmental Regulation,” Immunity 54, no. 10 (2021): 2305–2320.e11, 10.1016/j.immuni.2021.08.012.

33. A. Appios, J. Davies, S. Sirvent, et al., “Convergent Evolution of Monocyte Differentiation in Adult Skin Instructs Langerhans Cell Identity,” Science Immunology (2024), 10.1126/sciimmunol.adp0344.

34. J. D. Laman, P. J. M. Leenen, N. E. Annels, P. C. W. Hogendoorn, and R. M. Egeler, “Langerhans-Cell Histiocytosis ‘Insight into DC Biology,’” Trends in Immunology 24, no. 4 (2003): 190–196, 10.1016/s1471-4906(03)00063-2.

35. R. A. Schlegel, S. Krahling, M. K. Callahan, and P. Williamson, “CD14 Is a Component of Multiple Recognition Systems Used by Macrophages to Phagocytose Apoptotic Lymphocytes,” Cell Death & Differentiation 6, no. 6 (1999): 583–592, 10.1038/sj.cdd.4400529.

36. A. Maliah, N. Santana-Magal, S. Parikh, et al., “Crosslinking of Ly6a Metabolically Reprograms CD8 T Cells for Cancer Immunotherapy,” Nature Communications 15, no. 1 (2024): 8354, 10.1038/s41467-024-52079-x.

37. A. Fröhlich, S. Loick, E. G. Bawden, et al., “Comprehensive Analysis of Tumor Necrosis Factor Receptor TNFRSF9 (4-1BB) DNA Methylation with Regard to Molecular and Clinicopathological Features, Immune Infiltrates, and Response Prediction to Immunotherapy in Melanoma,” eBioMedicine 52 (2020), 10.1016/j.ebiom.2020.102647.

38. P. Stoitzner, C. H. Tripp, P. Douillard, S. Saeland, and N. Romani, “Migratory Langerhans Cells in Mouse Lymph Nodes in Steady State and Inflammation,” The Journal of Investigative Dermatology 125, no. 1 (2005): 116–125, 10.1111/j.0022-202X.2005.23757.x.

39. R. Tanaka, Y. Ichimura, N. Kubota, et al., “The Role of PD-L1 on Langerhans Cells in the Regulation of Psoriasis,” Journal of Investigative Dermatology 142, no. 12 (2022): 3167–3174.e9, 10.1016/j.jid.2022.06.006.

40. X. Wu, C. G. Briseño, V. Durai, et al., “Mafb Lineage Tracing to Distinguish Macrophages from Other Immune Lineages Reveals Dual Identity of Langerhans Cells,” The Journal of Experimental Medicine 213, no. 12 (2016): 2553–2565, 10.1084/jem.20160600.

41. R. Rauschmeier, C. Gustafsson, A. Reinhardt, et al., “Bhlhe40 and Bhlhe41 Transcription Factors Regulate Alveolar Macrophage Self-renewal and Identity,” The EMBO Journal 38, no. 19 (2019): e101233, 10.15252/embj.2018101233.

42. F. Jouenne, A. Benattia, and A. Tazi, “Mitogen-Activating Protein Kinase Pathway Alterations in Langerhans Cell Histiocytosis,” Current Opinion in Oncology 33, no. 2 (2021): 101–109, 10.1097/CCO.0000000000000707.

43. W. D. Shipman, S. Chyou, A. Ramanathan, et al., “A Protective Langerhans Cell-Keratinocyte Axis That Is Dysfunctional in Photosensitivity,” Science translational medicine 10, no. 454 (2018): eaap9527, 10.1126/scitranslmed.aap9527.

44. J. M. Kel, M. J. H. Girard-Madoux, B. Reizis, and B. E. Clausen, “TGF-Beta Is Required to Maintain the Pool of Immature Langerhans Cells in the Epidermis,” Journal of Immunology (Baltimore, Md.: 1950) 185, no. 6 (2010): 3248–3255, 10.4049/jimmunol.1000981.

45. M. R. Becker, Y. S. Choi, S. E. Millar, and M. C. Udey, “Wnt Signaling Influences the Development of Murine Epidermal Langerhans Cells,” The Journal of investigative dermatology 131, no. 9 (2011): 1861–1868, 10.1038/jid.2011.131.

46. S. A. Grace, A. M. Sutton, E. S. Armbrecht, C. I. Vidal, I. S. Rosman, and M. Y. Hurley, “P53 Is a Helpful Marker in Distinguishing Langerhans Cell Histiocytosis From Langerhans Cell Hyperplasia,” The American Journal of Dermatopathology 39, no. 10 (2017): 726–730, 10.1097/DAD.0000000000000778.

47. P. Novoszel, B. Drobits, M. Holcmann, et al., “The AP-1 Transcription Factors c-Jun and JunB Are Essential for CD8α Conventional Dendritic Cell Identity,” Cell Death & Differentiation 28, no. 8 (2021): 2404–2420, 10.1038/s41418-021-00765-4.

48. B. Kellersch, and T. Brocker, “Langerhans Cell Homeostasis in Mice Is Dependent on mTORC1 but Not mTORC2 Function,” Blood 121, no. 2 (2013): 298–307, 10.1182/blood-2012-06-439786.

49. N. Ma, E. Xu, Q. Luo, and G. Song, “Rac1: A Regulator of Cell Migration and a Potential Target for Cancer Therapy,” Molecules 28, no. 7 (2023): 2976, 10.3390/molecules28072976.

50. K. Seré, J.-H. Baek, J. Ober-Blöbaum, et al., “Two Distinct Types of Langerhans Cells Populate the Skin during Steady State and Inflammation,” Immunity 37, no. 5 (2012): 905–916, 10.1016/j.immuni.2012.07.019.

51. S. Kim, J. Ge, D. Kim, et al., “TXNIP-Mediated Crosstalk between Oxidative Stress and Glucose Metabolism,” PloS One 19, no. 2 (2024): e0292655, 10.1371/journal.pone.0292655.

52. G.-J. Zhao, K. Yin, Y. Fu, and C.-K. Tang, “The Interaction of ApoA-I and ABCA1 Triggers Signal Transduction Pathways to Mediate Efflux of Cellular Lipids,” Molecular Medicine 18, no. 2 (2012): 149–158, 10.2119/molmed.2011.00183.

53. F. Kemp, E. L. Braverman, and C. A. Byersdorfer, “Fatty Acid Oxidation in Immune Function,” Frontiers in Immunology 15 (2024): 1420336, 10.3389/fimmu.2024.1420336.

54. S. K. Wculek, I. Heras-Murillo, A. Mastrangelo, et al., “Oxidative Phosphorylation Selectively Orchestrates Tissue Macrophage Homeostasis,” Immunity 56, no. 3 (2023): 516–530.e9, 10.1016/j.immuni.2023.01.011.

55. S. M. Casillo, T. A. Gatesman, A. Chilukuri, et al., “An ERK5-PFKFB3 Axis Regulates Glycolysis and Represents a Therapeutic Vulnerability in Pediatric Diffuse Midline Glioma,” Cell Reports 43, no. 1 (2024): 113557, 10.1016/j.celrep.2023.113557.

56. S. Harshan, P. Dey, and S. Raghunathan, “Altered Transcriptional Regulation of Glycolysis in Circulating CD8+ T Cells of Rheumatoid Arthritis Patients,” Genes 13, no. 7 (2022): 1216, 10.3390/genes13071216.

57. S. Levi, M. Ripamonti, M. Dardi, A. Cozzi, and P. Santambrogio, “Mitochondrial Ferritin: Its Role in Physiological and Pathological Conditions,” Cells 10, no. 8 (2021): 1969, 10.3390/cells10081969.

58. D. E. Handy, E. Lubos, Y. Yang, et al., “Glutathione Peroxidase-1 Regulates Mitochondrial Function to Modulate Redox-Dependent Cellular Responses,” The Journal of Biological Chemistry 284, no. 18 (2009): 11913–11921, 10.1074/jbc.M900392200.

59. T. Mráček, Z. Drahota, and J. Houštěk, “The Function and the Role of the Mitochondrial Glycerol-3-Phosphate Dehydrogenase in Mammalian Tissues,” Biochimica et Biophysica Acta (BBA) - Bioenergetics 1827, no. 3 (2013): 401–410, 10.1016/j.bbabio.2012.11.014.

60. G. A. Gonzales, S. Huang, L. Wilkinson, et al., “The Pore-Forming Apolipoprotein APOL7C Drives Phagosomal Rupture and Antigen Cross-Presentation by Dendritic Cells,” Science Immunology 9, no. 101 (2024): eadn2168, 10.1126/sciimmunol.adn2168.

61. W. George Warren, M. Osborn, A. Yates, K. Wright, and S. E. O’Sullivan, “The Emerging Role of Fatty Acid Binding Protein 5 (FABP5) in Cancers,” Drug Discovery Today 28, no. 7 (2023): 103628, 10.1016/j.drudis.2023.103628.

62. V. Bolaños-Suárez, A. Alfaro, A. M. Espinosa, et al., “The mRNA and Protein Levels of the Glycolytic Enzymes Lactate Dehydrogenase A (LDHA) and Phosphofructokinase Platelet (PFKP) Are Good Predictors of Survival Time, Recurrence, and Risk of Death in Cervical Cancer Patients,” Cancer Medicine 12, no. 14 (2023): 15632–15649, 10.1002/cam4.6123.

63. H. Yu, L. Lin, Z. Zhang, H. Zhang, and H. Hu, “Targeting NF-κB Pathway for the Therapy of Diseases: Mechanism and Clinical Study,” Signal Transduction and Targeted Therapy 5, no. 1 (2020): 1–23, 10.1038/s41392-020-00312-6.

64. C. A. Dinarello, “Interleukin-1 in the Pathogenesis and Treatment of Inflammatory Diseases,” Blood 117, no. 14 (2011): 3720–3732, 10.1182/blood-2010-07-273417.

65. J. W. Larrick, V. Morhenn, Y. L. Chiang, and T. Shi, “Activated Langerhans Cells Release Tumor Necrosis Factor,” Journal of Leukocyte Biology 45, no. 5 (1989): 429–433, 10.1002/jlb.45.5.429.

66. S. Li, S. Han, K. Jin, et al., “SOCS2 Suppresses Inflammation and Apoptosis during NASH Progression through Limiting NF-κB Activation in Macrophages,” International Journal of Biological Sciences 17, no. 15 (2021): 4165–4175, 10.7150/ijbs.63889.

67. K. Ouwehand, S. W. Spiekstra, T. Waaijman, et al., “CCL5 and CCL20 Mediate Immigration of Langerhans Cells into the Epidermis of Full Thickness Human Skin Equivalents,” European Journal of Cell Biology 91, no. 10 (2012): 765–773, 10.1016/j.ejcb.2012.06.004.

68. O. Y. Elhussein Mohamed, A. Elazomi, M. S. Mohamed, and F. B. Abdalla, “Local Elevation of CCL22: A New Trend in Immunotherapy (Skin Model),” Journal of Cellular Immunotherapy 2, no. 2 (2016): 79–84, 10.1016/j.jocit.2015.12.001.

69. P. Conti, G. Ronconi, D. Lauritano, et al., “Impact of TNF and IL-33 Cytokines on Mast Cells in Neuroinflammation,” International Journal of Molecular Sciences 25, no. 6 (2024): 3248, 10.3390/ijms25063248.

70. J. D. Webster, and D. Vucic, “The Balance of TNF Mediated Pathways Regulates Inflammatory Cell Death Signaling in Healthy and Diseased Tissues,” Frontiers in Cell and Developmental Biology 8 (2020), 10.3389/fcell.2020.00365.

71. H. Tabata, H. Morita, H. Kaji, K. Tohyama, and Y. Tohyama, “Syk Facilitates Phagosome-Lysosome Fusion by Regulating Actin-Remodeling in Complement-Mediated Phagocytosis,” Scientific Reports 10, no. 1 (2020): 22086, 10.1038/s41598-020-79156-7.

72. V. B. Shah, T. R. Ozment-Skelton, D. L. Williams, and L. Keshvara, “Vav1 and PI3K Are Required for Phagocytosis of Beta-Glucan and Subsequent Superoxide Generation by Microglia,” Molecular Immunology 46, nos. 8–9 (2009): 1845–1853, 10.1016/j.molimm.2009.01.014.

73. A. Rousseau, and A. Bertolotti, “Regulation of Proteasome Assembly and Activity in Health and Disease,” Nature Reviews Molecular Cell Biology 19, no. 11 (2018): 697–712, 10.1038/s41580-018-0040-z.

74. D. A. Ferrington, and D. S. Gregerson, “Immunoproteasomes: Structure, Function, and Antigen Presentation,” Progress in molecular biology and translational science 109 (2012): 75–112, 10.1016/B978-0-12-397863-9.00003-1.

75. J. D. Mellor, M. P. Brown, H. R. Irving, J. R. Zalcberg, and A. Dobrovic, “A Critical Review of the Role of Fc Gamma Receptor Polymorphisms in the Response to Monoclonal Antibodies in Cancer,” Journal of Hematology & Oncology 6, no. 1 (2013): 1, 10.1186/1756-8722-6-1.

76. P. A. Roche, and K. Furuta, “The Ins and Outs of MHC Class II-Mediated Antigen Processing and Presentation,” Nature reviews. Immunology 15, no. 4 (2015): 203–216, 10.1038/nri3818.

77. N. M. Thielens, F. Tedesco, S. S. Bohlson, C. Gaboriaud, and A. J. Tenner, “C1q: A Fresh Look upon an Old Molecule,” Molecular immunology 89 (2017): 73–83, 10.1016/j.molimm.2017.05.025.

78. D. E. Hourcade, “The Role of Properdin in the Assembly of the Alternative Pathway C3 Convertases of Complement,” The Journal of Biological Chemistry 281, no. 4 (2006): 2128–2132, 10.1074/jbc.M508928200.

79. D. Ricklin, E. S. Reis, D. C. Mastellos, P. Gros, and J. D. Lambris, “Complement Component C3 - The ‘Swiss Army Knife’ of Innate Immunity and Host Defense,” Immunological reviews 274, no. 1 (2016): 33–58, 10.1111/imr.12500.

80. Y.-L. Wang, W.-W. Zhao, S.-M. Bai, et al., “MRNIP Condensates Promote DNA Double-Strand Break Sensing and End Resection,” Nature Communications 13, no. 1 (2022): 2638, 10.1038/s41467-022-30303-w.

81. D. Haensel, S. Jin, P. Sun, et al., “Defining Epidermal Basal Cell States during Skin Homeostasis and Wound Healing Using Single-Cell Transcriptomics,” Cell Reports 30, no. 11 (2020): 3932–3947.e6, 10.1016/j.celrep.2020.02.091.

82. S. Frech, and B. M. Lichtenberger, “Modulating Embryonic Signaling Pathways Paves the Way for Regeneration in Wound Healing,” Frontiers in Physiology 15 (2024): 1367425, 10.3389/fphys.2024.1367425.

83. J. M. Weiss, J. Sleeman, A. C. Renkl, et al., “An Essential Role for CD44 Variant Isoforms in Epidermal Langerhans Cell and Blood Dendritic Cell Function,” The Journal of Cell Biology 137, no. 5 (1997): 1137–1147.

84. A. A. Price, M. Cumberbatch, I. Kimber, and A. Ager, “Α6 Integrins Are Required for Langerhans Cell Migration from the Epidermis,” The Journal of Experimental Medicine 186, no. 10 (1997): 1725–1735.

85. K. Burke, and I. Gigli, “Receptors for Complement of Langerhans Cells,” The Journal of Investigative Dermatology 75, no. 1 (1980): 46–51, 10.1111/1523-1747.ep12521115.

86. R. Bai, Y. Guo, W. Liu, Y. Song, Z. Yu, and X. Ma, “The Roles of WNT Signaling Pathways in Skin Development and Mechanical-Stretch-Induced Skin Regeneration,” Biomolecules 13, no. 12 (2023): 1702, 10.3390/biom13121702.

87. N. S. Merle, R. Noe, L. Halbwachs-Mecarelli, V. Fremeaux-Bacchi, and L. T. Roumenina, “Complement System Part II: Role in Immunity,” Frontiers in Immunology 6 (2015), 10.3389/fimmu.2015.00257.

88. J. R. Dunkelberger, and W.-C. Song, “Complement and Its Role in Innate and Adaptive Immune Responses,” Cell Research 20, no. 1 (2010): 34–50, 10.1038/cr.2009.139.

89. T. Okada, H. Konishi, M. Ito, H. Kaneshima, and J. Asai, “Developmental Expression of C3 Receptor on Murine Epidermal Langerhans Cells during Ontogeny,” Archives of Dermatological Research 280, no. 1 (1988): 39–44, 10.1007/BF00412687.

90. P. Hillmen, J. Szer, I. Weitz, et al., “Pegcetacoplan versus Eculizumab in Paroxysmal Nocturnal Hemoglobinuria,” New England Journal of Medicine 384, no. 11 (2021): 1028–1037, 10.1056/NEJMoa2029073.

91. Y. Liu, A. Beyer, and R. Aebersold, “On the Dependency of Cellular Protein Levels on mRNA Abundance,” Cell 165, no. 3 (2016): 535–550, 10.1016/j.cell.2016.03.014.

92. C. Vogel, and E. M. Marcotte, “Insights into the Regulation of Protein Abundance from Proteomic and Transcriptomic Analyses,” Nature Reviews Genetics 13, no. 4 (2012): 227–232, 10.1038/nrg3185.

93. M. D. Luecken, and F. J. Theis, “Current Best Practices in Single-Cell RNA-Seq Analysis: A Tutorial,” Molecular Systems Biology 15, no. 6 (2019): e8746, 10.15252/msb.20188746.

94. T. Stuart, and R. Satija, “Integrative Single-Cell Analysis,” Nature Reviews. Genetics 20, no. 5 (2019): 257–272, 10.1038/s41576-019-0093-7.

95. A. Kapp, and E. Schöpf, “Involvement of Complement in Atopic Dermatitis,” Acta Dermato-Venereologica. Supplementum 114 (1985): 152–154, 10.2340/00015555114152154.

96. E. W. Rosenberg, P. W. Noah, R. J. Wyatt, R. M. Jones, and W. P. Kolb, “Complement Activation in Psoriasis,” Clinical and Experimental Dermatology 15, no. 1 (1990): 16–20, 10.1111/j.1365-2230.1990.tb02011.x.

97. J. Panelius, and S. Meri, “Complement System in Dermatological Diseases – Fire Under the Skin,” Frontiers in Medicine 2 (2015): 3, 10.3389/fmed.2015.00003.

98. L. Dang, L. He, Y. Wang, J. Xiong, B. Bai, and Y. Li, “Role of the Complement Anaphylatoxin C5a-Receptor Pathway in Atopic Dermatitis in Mice,” Molecular Medicine Reports 11, no. 6 (2015): 4183–4189, 10.3892/mmr.2015.3301.

99. Q.-Y. Zheng, S.-J. Liang, F. Xu, et al., “C5a/C5aR1 Pathway Is Critical for the Pathogenesis of Psoriasis,” Frontiers in Immunology 10 (2019): 1866, 10.3389/fimmu.2019.01866.

100. Q. Zheng, F. Xu, Y. Yang, et al., “C5a/C5aR1 Mediates IMQ-Induced Psoriasiform Skin Inflammation by Promoting IL-17A Production from Γδ-T Cells,” The FASEB Journal 34, no. 8 (2020): 10590–10604, 10.1096/fj.202000384R.

101. P. Rompolas, E. R. Deschene, G. Zito, et al., “Live Imaging of Stem Cell and Progeny Behaviour in Physiological Hair-Follicle Regeneration,” Nature 487, no. 7408 (2012): 496–499, 10.1038/nature11218.

102. A. Butler, P. Hoffman, P. Smibert, E. Papalexi, and R. Satija, “Integrating Single-Cell Transcriptomic Data across Different Conditions, Technologies, and Species,” Nature Biotechnology 36, no. 5 (2018): 411–420, 10.1038/nbt.4096.

103. F. A. Wolf, P. Angerer, and F. J. Theis, “SCANPY: Large-Scale Single-Cell Gene Expression Data Analysis,” Genome Biology 19, no. 1 (2018): 15, 10.1186/s13059-017-1382-0.

104. Samuel Marsh, Maëlle Salmon, Paul Hoffman, kew24, and Mustafa Samet Pir, “Samuel-Marsh/scCustomize: Release 3.2.4,” samuel-marsh/scCustomize: Release 3.2.4, Zenodo 2025, 10.5281/ZENODO.5706430.

105. J. A. Bibby, D. Agarwal, T. Freiwald, et al., “Systematic Single-Cell Pathway Analysis to Characterize Early T Cell Activation,” Cell Reports 41, no. 8 (2022): 111697, 10.1016/j.celrep.2022.111697.

106. A. Liberzon, C. Birger, H. Thorvaldsdóttir, M. Ghandi, J. P. Mesirov, and P. Tamayo, “The Molecular Signatures Database Hallmark Gene Set Collection,” Cell Systems 1, no. 6 (2015): 417–425, 10.1016/j.cels.2015.12.004.

107. A. Subramanian, P. Tamayo, V. K. Mootha, et al., “Gene Set Enrichment Analysis: A Knowledge-Based Approach for Interpreting Genome-Wide Expression Profiles,” Proceedings of the National Academy of Sciences 102, no. 43 (2005): 15545–15550, 10.1073/pnas.0506580102.

108. M. Milacic, D. Beavers, P. Conley, et al., “The Reactome Pathway Knowledgebase 2024,” Nucleic Acids Research 52, no. D1 (2024): D672–D678, 10.1093/nar/gkad1025.

109. A. Agrawal, H. Balcı, K. Hanspers, et al., “WikiPathways 2024: Next Generation Pathway Database,” Nucleic Acids Research 52, no. D1 (2024): D679–D689, 10.1093/nar/gkad960.

110. S. Aibar, C. B. González-Blas, T. Moerman, et al., “SCENIC: Single-Cell Regulatory Network Inference and Clustering,” Nature Methods 14, no. 11 (2017): 1083–1086, 10.1038/nmeth.4463.

111. G. La Manno, R. Soldatov, A. Zeisel, et al., “RNA Velocity of Single Cells,” Nature 560, no. 7719 (2018): 494–498, 10.1038/s41586-018-0414-6.

112. V. Bergen, M. Lange, S. Peidli, F. A. Wolf, and F. J. Theis, “Generalizing RNA Velocity to Transient Cell States through Dynamical Modeling,” Nature Biotechnology 38, no. 12 (2020): 1408–1414, 10.1038/s41587-020-0591-3.

113. K. Street, D. Risso, R. B. Fletcher, et al., “Slingshot: Cell Lineage and Pseudotime Inference for Single-Cell Transcriptomics,” BMC genomics 19, no. 1 (2018): 477, 10.1186/s12864-018-4772-0.

114. K. Van den Berge, H. Roux de Bézieux, K. Street, et al., “Trajectory-Based Differential Expression Analysis for Single-Cell Sequencing Data,” Nature Communications 11, no. 1 (2020): 1201, 10.1038/s41467-020-14766-3.

115. M. Kang, G. S. Gulati, E. L. Brown, et al., “Improved Reconstruction of Single-Cell Developmental Potential with CytoTRACE 2,” Nature Methods 22, no. 11 (2025): 2258–2263, 10.1038/s41592-025-02857-2.

116. D. Dimitrov, P. S. L. Schäfer, E. Farr, et al., “LIANA+ Provides an All-in-One Framework for Cell–Cell Communication Inference,” Nature Cell Biology 26, no. 9 (2024): 1613–1622, 10.1038/s41556-024-01469-w.

117. J. Schindelin, I. Arganda-Carreras, E. Frise, et al., “Fiji: An Open-Source Platform for Biological-Image Analysis,” Nature Methods 9, no. 7 (2012): 676–682, 10.1038/nmeth.2019.

118. C. Stringer, T. Wang, M. Michaelos, and M. Pachitariu, “Cellpose: A Generalist Algorithm for Cellular Segmentation,” Nature Methods 18, no. 1 (2021): 100–106, 10.1038/s41592-020-01018-x.

119. L. Mac-Daniel, M. R. Buckwalter, P. Gueirard, and R. Ménard, “Myeloid Cell Isolation from Mouse Skin and Draining Lymph Node Following Intradermal Immunization with Live Attenuated Plasmodium Sporozoites,” Journal of Visualized Experiments: JoVE no. 111 (2016): 53796, 10.3791/53796.

120. A. J. Schmidt, G. D. Wright, F. Ronchese, and K. M. Price, “Skin Whole-Mount Immunofluorescent Staining Protocol, 3D Visualization, and Spatial Image Analysis,” 10.1002/cpz1.820.

