## Supporting Information for "A Transcriptionally Distinct Intermediate Activation State Precedes Langerhans Cell Migration from the Epidermis"

**Figure S1. Quality control and preprocessing define a robust single-cell RNA sequencing (scRNA-seq) dataset for downstream analysis of Langerhans cell (LC) states. A.** Diagram of the scRNA-seq workflow from LC isolation to downstream analysis using Seurat. **B.** Barcode rank plots generated using the Barcode_Plot function from the scCustomize package, showing ranked barcodes by total unique molecular identifier (UMI) counts for each sample. Inflection and knee points are indicated. **C.** Quality control metrics before and after filtering, including nFeature_RNA (number of detected genes per cell), nCount_RNA (total UMI counts per cell), percent_mito (percentage of mitochondrial transcripts), and percent_ribo (percentage of ribosomal transcripts). **D.** Uniform Manifold Approximation and Projection (UMAP) visualizations of the scRNA-seq dataset at different preprocessing stages. Left: unfiltered dataset. Middle: filtered dataset annotated into three LC clusters. Right: filtered dataset annotated at higher clustering resolution into eight clusters (for **Figure 3A, C**).

**Figure S2. Time-course analysis defines 46 h as an early response time point for studying Langerhans cell (LC) spatial reorganization. A.** Representative intravital images of LCs repeatedly acquired from the same epidermal area at the indicated time points following croton oil treatment (left), with corresponding Voronoi diagrams generated from the segmented LC positions (right) (n = 3 mice). **B.** Representative quantification of the LC spatial distribution based on the Voronoi diagram (n = 3 mice). **C.** Representative quantification of the minimum intercellular distance between LCs, calculated as the average distance to the three nearest neighboring LCs (***p < 0.001, ****p < 0.0001; n = 3 mice). **D.** Representative quantification of LC numbers over time following croton oil treatment (n = 3 mice).

**Figure S3. Flow cytometry gating strategy for consistent identification of activated Langerhans, T, and B cells across conditions.** **A.** Gating of epidermal Langerhans cells (LCs): cells, singlets, live cells, and Cd207-eGFP⁺ Krt14-mCherry⁻ LCs. **B.** Marker versus CXCR4 expression within LCs for CD137 (Tnfrsf9), Ly-6A/E (Sca-1), CD14, and CD80 in control and croton/acetone-treated mice. **C.** Gating of draining lymph node cells: CD3⁺ and CD4⁺ T cells, CD69⁺ cells within CD3⁺ and CD4⁺ T cells, and B220⁺ B cells.

**Figure S4. Quantitative flow cytometry analysis of LC activation markers and lymph node immune cells across conditions.** Quantification of control, croton/acetone-, and croton/acetone + ovalbumin (OVA)-treated mice, gated as in Figure S3. **A–C.** Percentage of ear epidermal LCs positive for CD137 (Tnfrsf9), Ly-6A/E (Sca-1), and CD14. **D–F.** Marker⁺ LCs from A–C subdivided into CXCR4⁻ (filled) and CXCR4⁺ (open) fractions. **G, H.** Percentage of LCs positive for CD80 and CD86. **I, J.** Percentage of CD69⁺ cells within CD3⁺ and CD4⁺ T cells in the draining lymph node. **K.** Percentage of B220⁺ cells among live draining lymph node cells. Each dot represents one mouse; bars show mean ± SD; n = 4 mice per group, except n = 3 for control in I. A–C and G–K: two-tailed Welch’s t test versus control. D–F: two-tailed paired t test of CXCR4⁻ versus CXCR4⁺ within each treatment. P values were adjusted using the Holm method for the two comparisons in each panel.

**Figure S5. Expression of skin cell-type markers across Langerhans cell (LC) states supports their distinct transcriptional identity.** Dot plot showing the relative expression of selected marker genes associated with mouse dermal and epidermal cell types across steady, activated, and migratory LC states.

**Figure S6. Pseudotime analysis reveals dynamic gene expression changes along the Langerhans cell (LC) activation trajectory.** Heatmap showing the smoothed expression patterns of the top 150 genes significantly associated with pseudotime. Gene expression was modeled using tradeSeq, smoothed across a pseudotime grid, and scaled via a gene-wise z-score. Columns represent ordered pseudotime progression and rows represent hierarchically clustered genes.

**Figure S7. Differential gene expression defines distinct transcriptional identities of Langerhans cell (LC) states.** Heatmap showing the expression of the top differentially expressed genes across clusters in the Seurat object. Marker genes were identified using Seurat FindAllMarkers (Wilcoxon rank-sum test; only pos = TRUE, logfc. threshold = 0.25, min.pct = 0.40, return threshold = 0.05). For each cluster, the top 50 marker genes ranked according to average log2 fold change (avg_log2FC) were selected and combined into a non-redundant gene set (n = 50 genes per cluster) for visualization.

**Figure S8. Migratory, activated, and steady Langerhans cell (LC) states differ in cell-cycle phase distribution.** Distribution of cell-cycle phases across the three main LC clusters (steady, activated, and migratory). Migratory LCs were enriched in the G1 phase and depleted in the S phase, whereas steady-state LCs had a higher proportion of cells in the S phase. These differences indicate variations in cell-cycle composition across LC states.

**Figure S9. Spatial distribution of activated Langerhans cells (LC) following skin inflammation and injury.** **A.** Representative whole-mount staining images of epidermis under homeostatic conditions and two days after croton oil treatment. CD137 is shown in green, CD207 in red, and 4′,6-diamidino-2-phenylindole (DAPI) in blue. Arrows indicate CD137^+^ LCs (n = 3 mice). **B.** Quantification of CD207^+^ CD137^+^ double-positive LCs in homeostatic and croton oil-treated skin (n = 3 mice). **C.** Representative whole-mount staining images of epidermal skin two days after punch biopsy. CD137 is shown in green, CD207 in red, and DAPI in blue. Arrows indicate CD137^+^ LCs. (n = 3 mice). **D.** Quantification of CD207^+^ CD137^+^ double-positive LCs in regions adjacent to the wound and regions distant from the wound (n = 3 mice).

**Figure S10. CytoTRACE2 analysis reveals a gradient of cellular potency within homeostatic Langerhans cells (LC).** **A.** Uniform Manifold Approximation and Projection (UMAP) visualization of the untreated control sample was analyzed independently, showing the clustering of LC populations. **B.** CytoTRACE2 analysis revealed a gradient of cellular potency across LCs, with the predicted differentiation potential inferred from the transcriptional diversity and displayed upon UMAP embedding.

**Figure S11. *Pcdh7* contributes to the spatial organization of Langerhans cells (LC) in the epidermis. A.** Whole-mount Langerin staining of the epidermis from control mice showing spatial organization of LCs under homeostatic conditions (n = 3 mice). **B.** Whole-mount Langerin staining of the epidermis from *Pcdh7* knockout mice, revealing altered LC spatial organization (n = 3 mice). **C.** Quantification of LC spatial distribution demonstrates changes in LC patterning in the *Pcdh7*-deficient epidermis compared to controls (n = 3 mice). Scale bar, 50 µm.

**Figure S12. *C3* and *C5 (Hc)* expression across skin cell populations in wounded and unwounded skin.** Dot plots show *C3* (left) and *C5* (right) expression across major skin cell populations from the integrated Haensel *et al*. (2020) dataset under unwounded and wounded conditions. Dot size represents the proportion of cells expressing each gene, and color intensity indicates scaled expression levels.

**Figure S13. Graphical model of complement-mediated interactions between Langerhans cells (LC) and fibroblasts during epidermal immune responses.**

Graphical model summarizing complement-related gene expression and predicted interactions between LCs and fibroblasts in the epidermis. Activated LCs express C1q subunits, whereas fibroblast populations, including wingless-related integration site (WNT)-modulated fibroblasts, express components required for complement activation, such as C1r, C1s, C3, and C4b. These predicted interactions suggest potential complement-related coordination between LCs and fibroblasts following tissue injury.

**Video S1. Intravital time-lapse imaging of Langerhans cell (LC) behavior 48 h after croton oil/acetone treatment.** Mouse ears were treated with croton oil/acetone, and intravital time-lapse imaging was performed 48 h after treatment. An 8-h recording captures the dynamic behavior of LCs within the epidermis following skin stimulation.

**Table S1. Differentially expressed marker genes across steady, activated, and migratory Langerhans cell (LC) states.** Marker genes were identified using the Wilcoxon rank-sum test to compare gene expression between the three stages. Displayed genes meet the following thresholds: average log2 fold-change (avg_log2FC) > 0.585, Benjamini–Hochberg adjusted p-value (p-val_adj) < 0.05, and expression detected in at least 10% of cells (pct ≥ 0.1). Only the positive markers enriched at each stage relative to the others were included.

**Table S2. Differentially expressed marker genes of homeostatic Langerhans cell (LC) populations in the control sample.** This table lists the positive marker genes identified using the Wilcoxon rank-sum test for homeostatic LC populations in the control samples. Only genes with an average log2 fold change > 0.585, adjusted p-value < 0.05, and expression in at least 10% of cells were included.

**Table S3. Predicted ligand–receptor interactions between Langerhans cells (LC) and skin cell populations.** Table summarizing predicted ligand–receptor interactions inferred using LIANA++ following integration of the study LC dataset with Haensel *et al*. (2020). The table includes source and target cell populations, ligand–receptor pairs, and associated interaction metrics (expression levels, statistical scores, and ranking values).
