## Supplemental Figures for "A Transcriptionally Distinct Intermediate Activation State Precedes Langerhans Cell Migration from the Epidermis"

### Figure S1

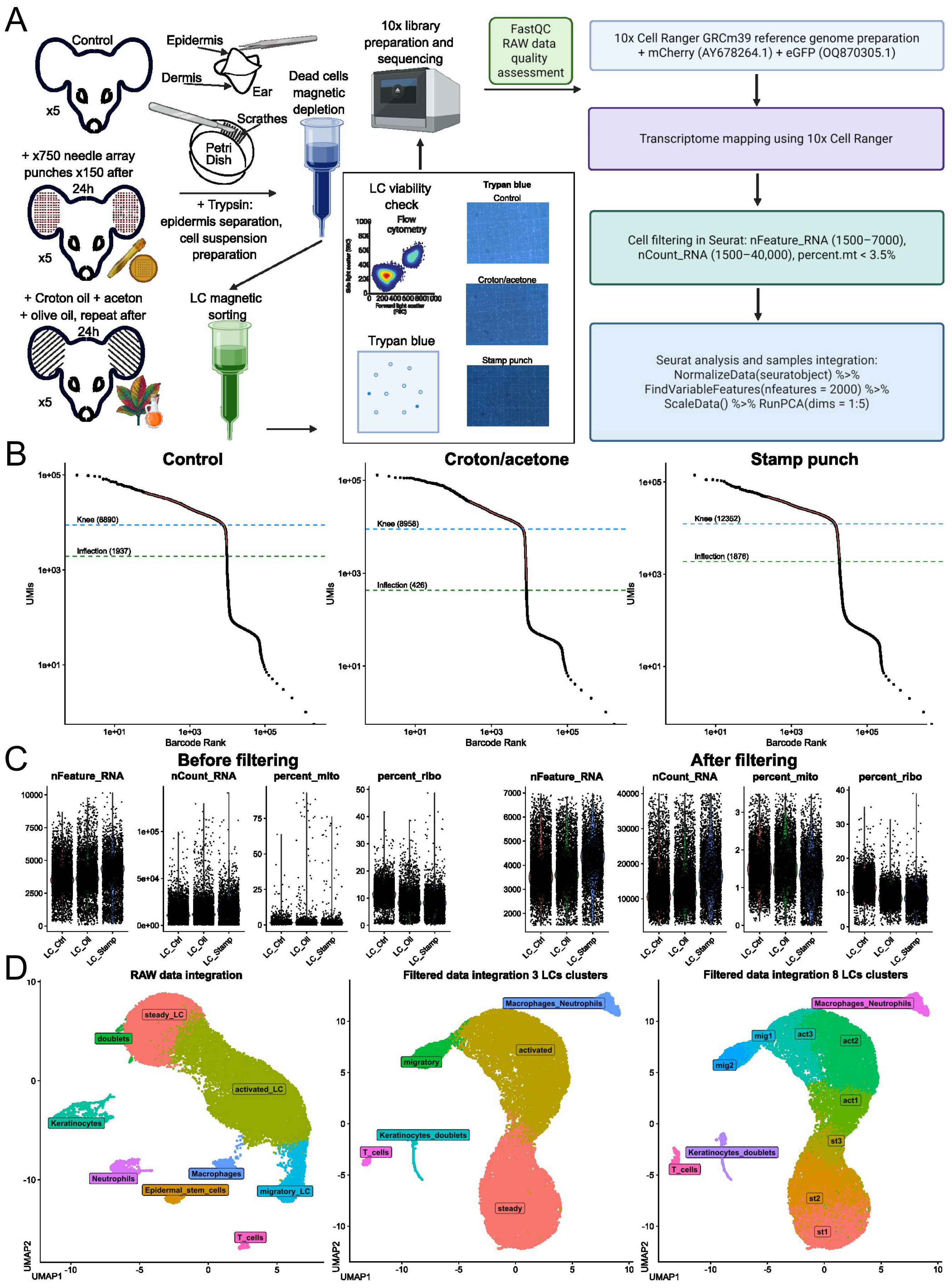

A
Control
x5
Epidermis
Dermis
Ear
Scrathes
Petri Dish
+ x750 needle array punches x150 after 24h
x5
+ Trypsin: epidermis separation, cell suspension preparation
Dead cells magnetic depletion
+ Croton oil + aceton + olive oil, repeat after 24h
x5
LC magnetic sorting
10x library preparation and sequencing
FastQC RAW data quality assessment
LC viability check
Flow cytometry
Trypan blue
Trypan blue
Control
Croton/acetone
Stamp punch
10x Cell Ranger GRCm39 reference genome preparation + mCherry (AY678264.1) + eGFP (OQ870305.1)
Transcriptome mapping using 10x Cell Ranger
Cell filtering in Seurat: nFeature_RNA (1500–7000), nCount_RNA (1500–40,000), percent.mt < 3.5%
Seurat analysis and samples integration: NormalizeData(seuratobject) %>% FindVariableFeatures(nfeatures = 2000) %>% ScaleData() %>% RunPCA(dims = 1:5)
B
Control
Croton/acetone
Stamp punch
Knee (8890)
Inflection (1937)
Knee (8958)
Inflection (426)
Knee (12352)
Inflection (1876)
UMIs
Barcode Rank
C
Before filtering
After filtering
nFeature_RNA
nCount_RNA
percent_mito
percent_ribo
LC_Ctrl
LC_Oil
LC_Stamp
D
RAW data integration
Filtered data integration 3 LCs clusters
Filtered data integration 8 LCs clusters
steady_LC
doublets
activated_LC
Keratinocytes
Neutrophils
Macrophages
Epidermal_stem_cells
migratory_LC
T_cells
Macrophages_Neutrophils
activated
migratory
Keratinocytes_doublets
steady
mig1
mig2
act1
act2
act3
st1
st2
st3
UMAP1
UMAP2

### Figure S2

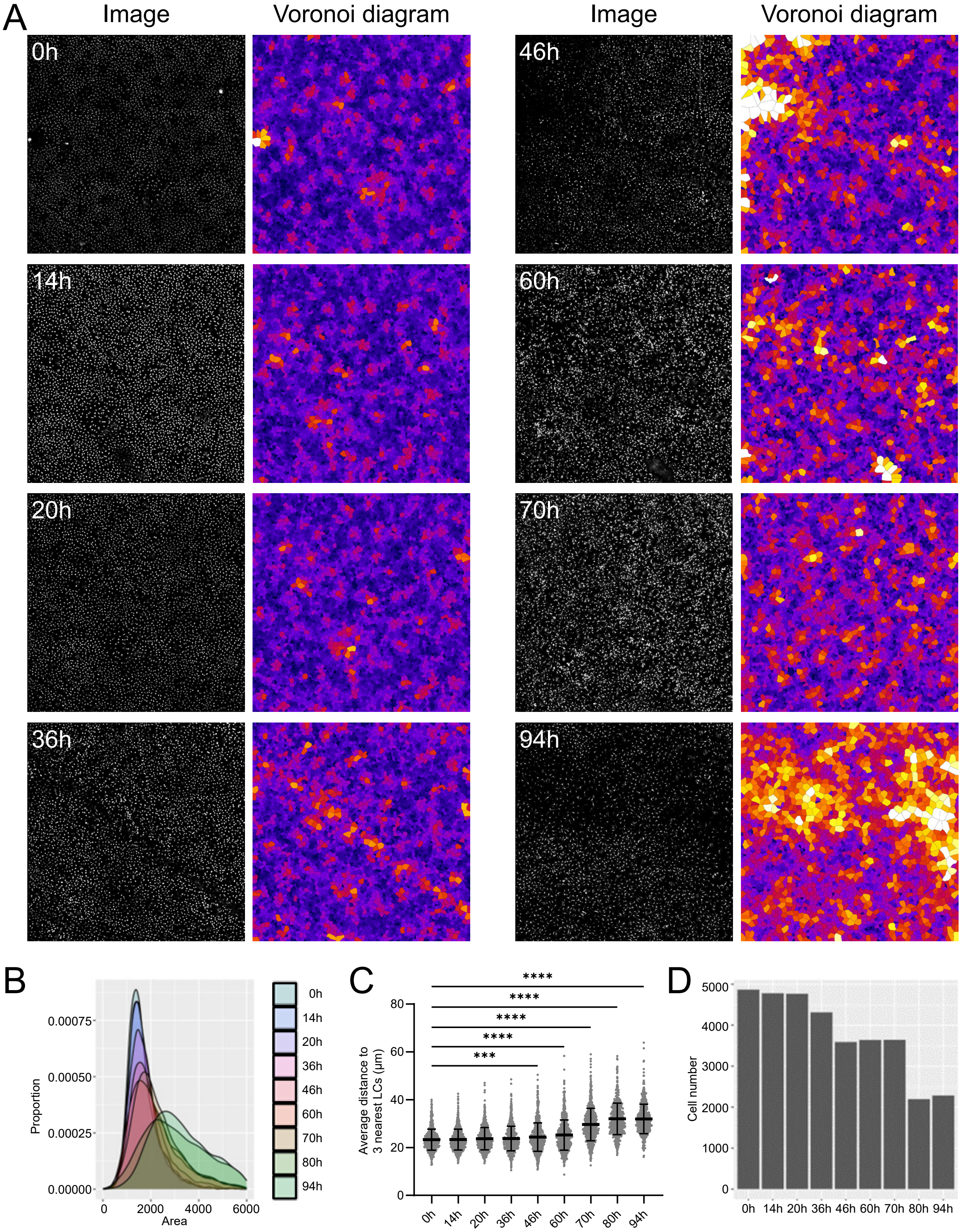
A
Image
Voronoi diagram
Image
Voronoi diagram
0h
14h
20h
36h
46h
60h
70h
94h
B
Proportion
0.00075
0.00050
0.00025
0.00000
0
2000
4000
6000
Area
0h
14h
20h
36h
46h
60h
70h
80h
94h
C
Average distance to 3 nearest LCs (μm)
****
****
****
****
***
80
60
40
20
0
0h 14h 20h 36h 46h 60h 70h 80h 94h
D
Cell number
5000
4000
3000
2000
1000
0
0h 14h 20h 36h 46h 60h 70h 80h 94h

### Figure S3

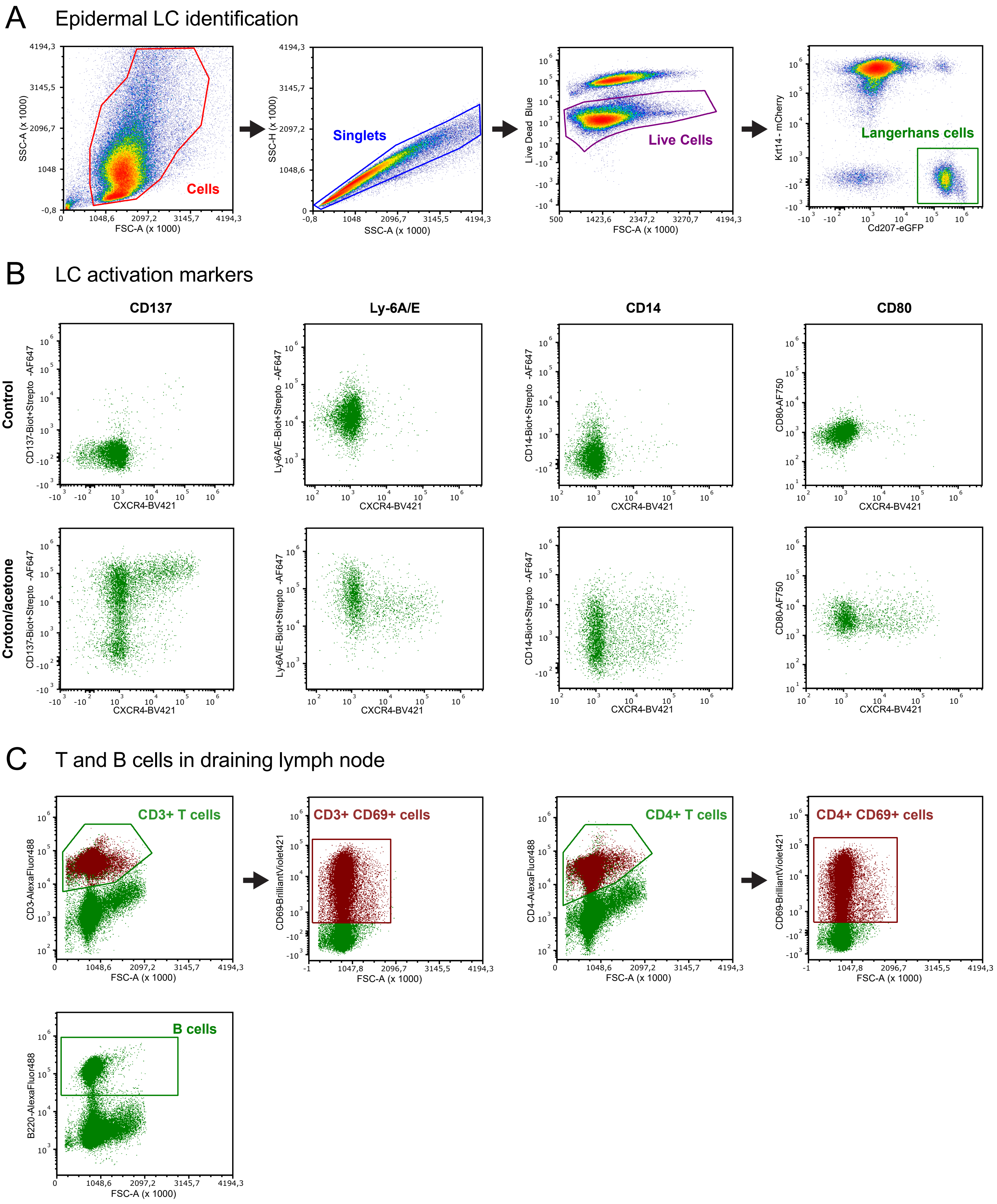

A
Epidermal LC identification
SSC-A (x 1000)
FSC-A (x 1000)
Cells
SSC-H (x 1000)
SSC-A (x 1000)
Singlets
Live Dead Blue
FSC-A (x 1000)
Live Cells
Krt14 - mCherry
Cd207-eGFP
Langerhans cells
B
LC activation markers
CD137
Ly-6A/E
CD14
CD80
Control
Croton/acetone
CD137-Biot+Strepto -AF647
Ly-6A/E-Biot+Strepto -AF647
CD14-Biot+Strepto -AF647
CD80-AF750
CXCR4-BV421
C
T and B cells in draining lymph node
CD3+ T cells
CD3+ CD69+ cells
CD4+ T cells
CD4+ CD69+ cells
B cells
CD3-AlexaFluor488
CD69-BrilliantViolet421
CD4-AlexaFluor488
B220-AlexaFluor488
FSC-A (x 1000)

### Figure S4

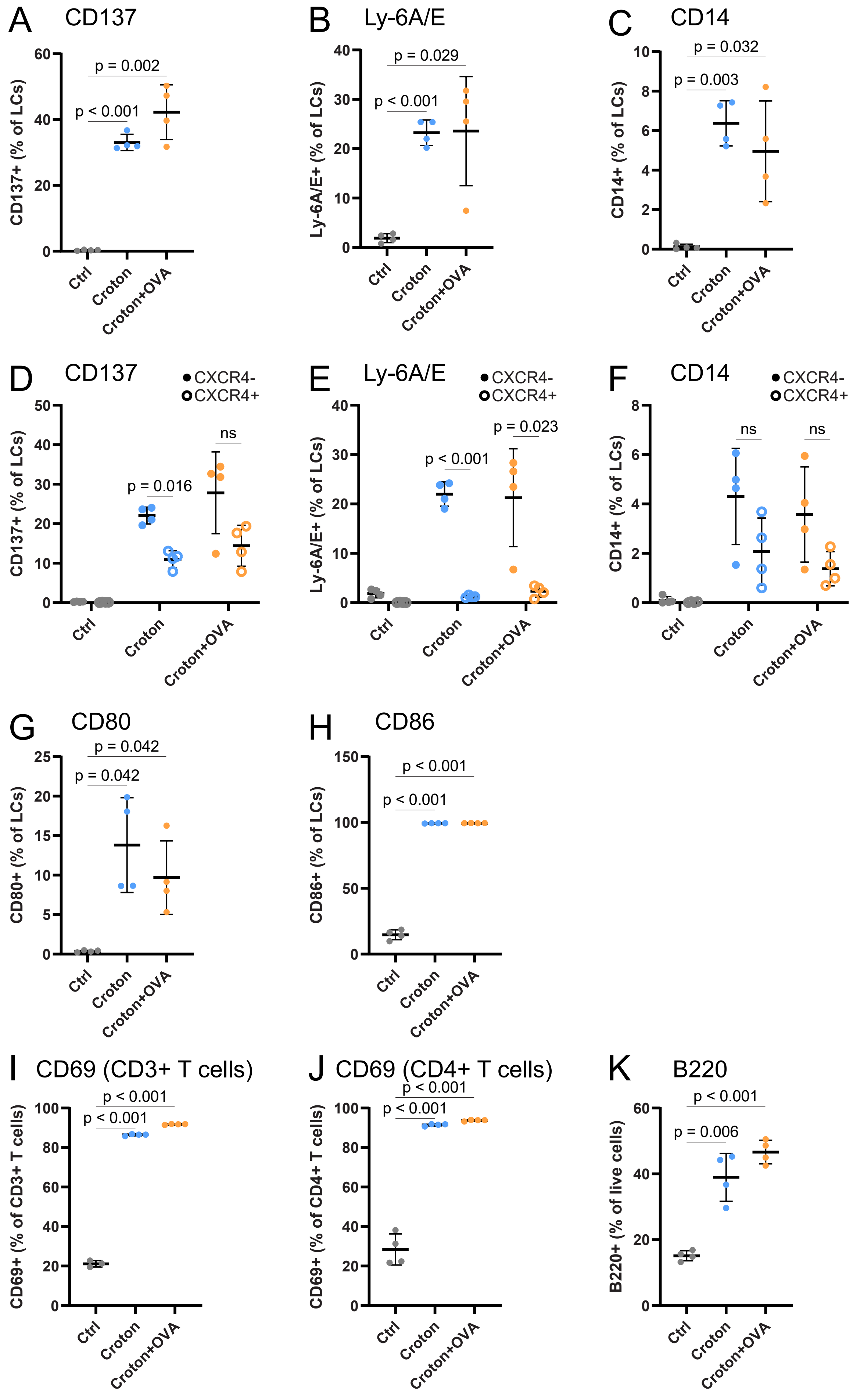
A
CD137
CD137+ (% of LCs)
p < 0.001
p = 0.002
B
Ly-6A/E
Ly-6A/E+ (% of LCs)
p < 0.001
p = 0.029
C
CD14
CD14+ (% of LCs)
p = 0.003
p = 0.032
D
CD137
CXCR4-
CXCR4+
p = 0.016
ns
E
Ly-6A/E
p < 0.001
p = 0.023
F
CD14
ns
ns
G
CD80
CD80+ (% of LCs)
p = 0.042
p = 0.042
H
CD86
CD86+ (% of LCs)
p < 0.001
p < 0.001
I
CD69 (CD3+ T cells)
CD69+ (% of CD3+ T cells)
p < 0.001
p < 0.001
J
CD69 (CD4+ T cells)
CD69+ (% of CD4+ T cells)
p < 0.001
p < 0.001
K
B220
B220+ (% of live cells)
p = 0.006
p < 0.001
Ctrl
Croton
Croton+OVA

### Figure S5

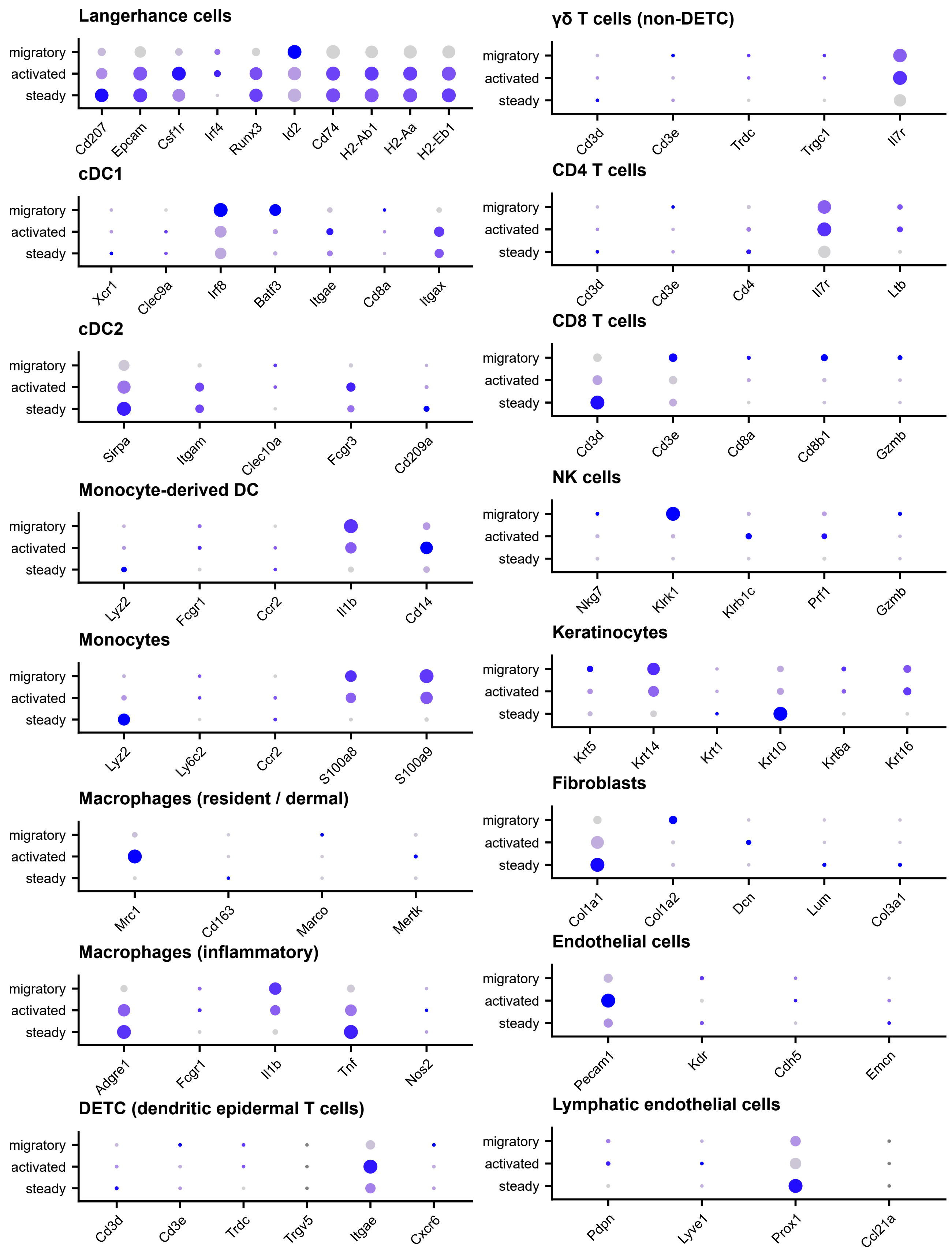

Langerhance cells
migratory
activated
steady
Cd207
Epcam
Csf1r
Irf4
Runx3
Id2
Cd74
H2-Ab1
H2-Aa
H2-Eb1
cDC1
Xcr1
Clec9a
Irf8
Batf3
Itgae
Cd8a
Itgax
cDC2
Sirpa
Itgam
Clec10a
Fcgr3
Cd209a
Monocyte-derived DC
Lyz2
Fcgr1
Ccr2
Il1b
Cd14
Monocytes
Lyz2
Ly6c2
Ccr2
S100a8
S100a9
Macrophages (resident / dermal)
Mrc1
Cd163
Marco
Mertk
Macrophages (inflammatory)
Adgre1
Fcgr1
Il1b
Tnf
Nos2
DETC (dendritic epidermal T cells)
Cd3d
Cd3e
Trdc
Trgv5
Itgae
Cxcr6
γδ T cells (non-DETC)
Cd3d
Cd3e
Trdc
Trgc1
Il7r
CD4 T cells
Cd3d
Cd3e
Cd4
Il7r
Ltb
CD8 T cells
Cd3d
Cd3e
Cd8a
Cd8b1
Gzmb
NK cells
Nkg7
Klrk1
Klrb1c
Prf1
Gzmb
Keratinocytes
Krt5
Krt14
Krt1
Krt10
Krt6a
Krt16
Fibroblasts
Col1a1
Col1a2
Dcn
Lum
Col3a1
Endothelial cells
Pecam1
Kdr
Cdh5
Emcn
Lymphatic endothelial cells
Pdpn
Lyve1
Prox1
Ccl21a

### Figure S6

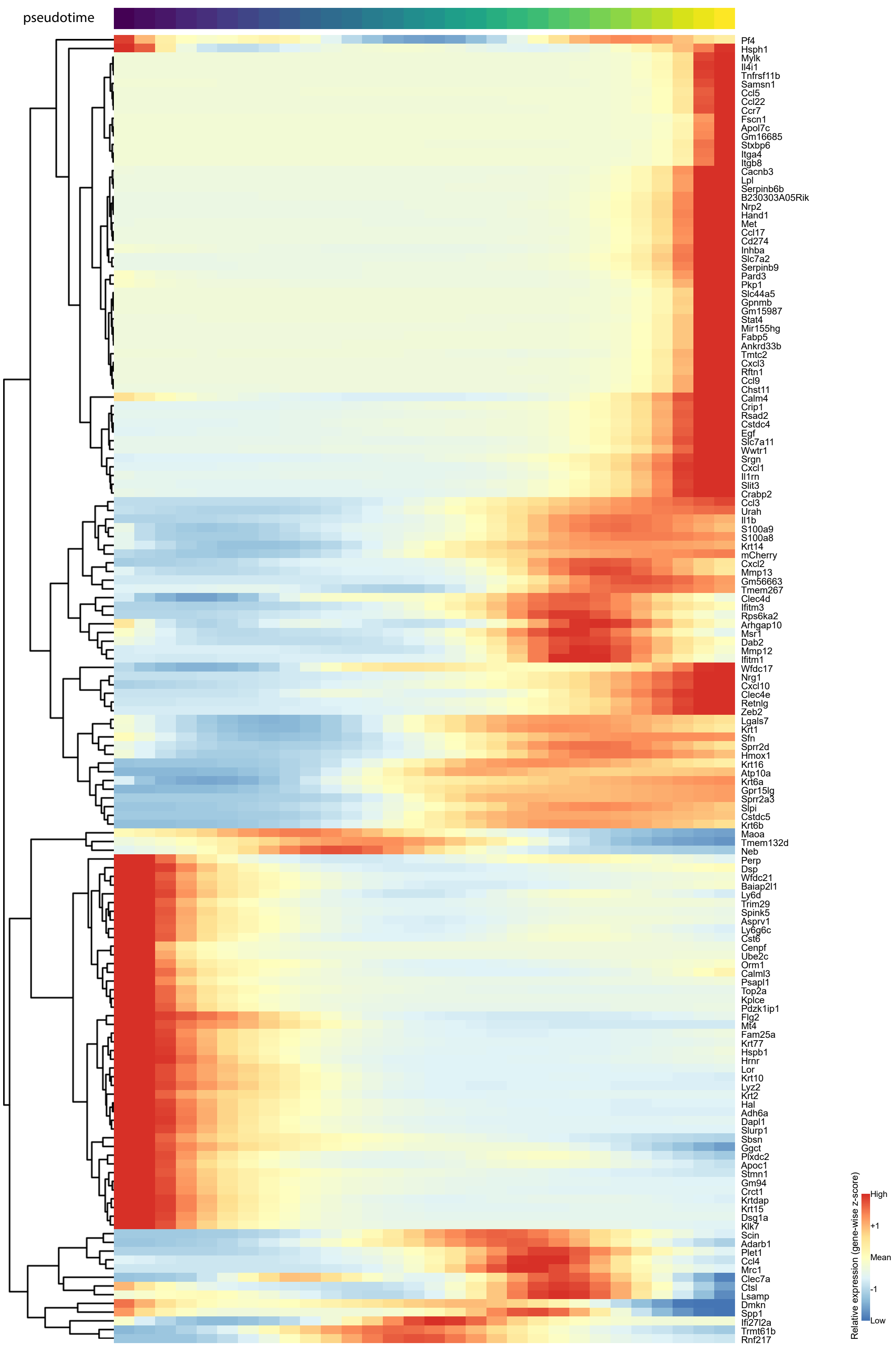
pseudotime
Pf4
Hsph1
Mylk
Il4i1
Tnfrsf11b
Samsn1
Ccl5
Ccl22
Ccr7
Fscn1
Apol7c
Gm16685
Stxbp6
Itga4
Itgb8
Cacnb3
Lpl
Serpinb6b
B230303A05Rik
Nrp2
Hand1
Met
Ccl17
Cd274
Inhba
Slc7a2
Serpinb9
Pard3
Pkp1
Slc44a5
Gpnmb
Gm15987
Stat4
Mir155hg
Fabp5
Ankrd33b
Tmtc2
Cxcl3
Rftn1
Ccl9
Chst11
Calm4
Crip1
Rsad2
Cstdc4
Egf
Slc7a11
Wwtr1
Srgn
Cxcl1
Il1rn
Slit3
Crabp2
Ccl3
Urah
Il1b
S100a9
S100a8
Krt14
mCherry
Cxcl2
Mmp13
Gm56663
Tmem267
Clec4d
Ifitm3
Rps6ka2
Arhgap10
Msr1
Dab2
Mmp12
Ifitm1
Wfdc17
Nrg1
Cxcl10
Clec4e
Retnlg
Zeb2
Lgals7
Krt1
Sfn
Sprr2d
Hmox1
Krt16
Atp10a
Krt6a
Gpr15lg
Sprr2a3
Slpi
Cstdc5
Krt6b
Maoa
Tmem132d
Neb
Perp
Dsp
Wfdc21
Baiap2l1
Ly6d
Trim29
Spink5
Asprv1
Ly6g6c
Cst6
Cenpf
Ube2c
Orm1
Calml3
Psapl1
Top2a
Kplce
Pdzk1ip1
Flg2
Mt4
Fam25a
Krt77
Hspb1
Hrnr
Lor
Krt10
Lyz2
Krt2
Hal
Adh6a
Dapl1
Slurp1
Sbsn
Ggct
Plxdc2
Apoc1
Stmn1
Gm94
Crct1
Krtdap
Krt15
Dsg1a
Klk7
Scin
Adarb1
Plet1
Ccl4
Mrc1
Clec7a
Ctsl
Lsamp
Dmkn
Spp1
Ifi27l2a
Trmt61b
Rnf217
Relative expression (gene-wise z-score)
High
+1
Mean
-1
Low

### Figure S7

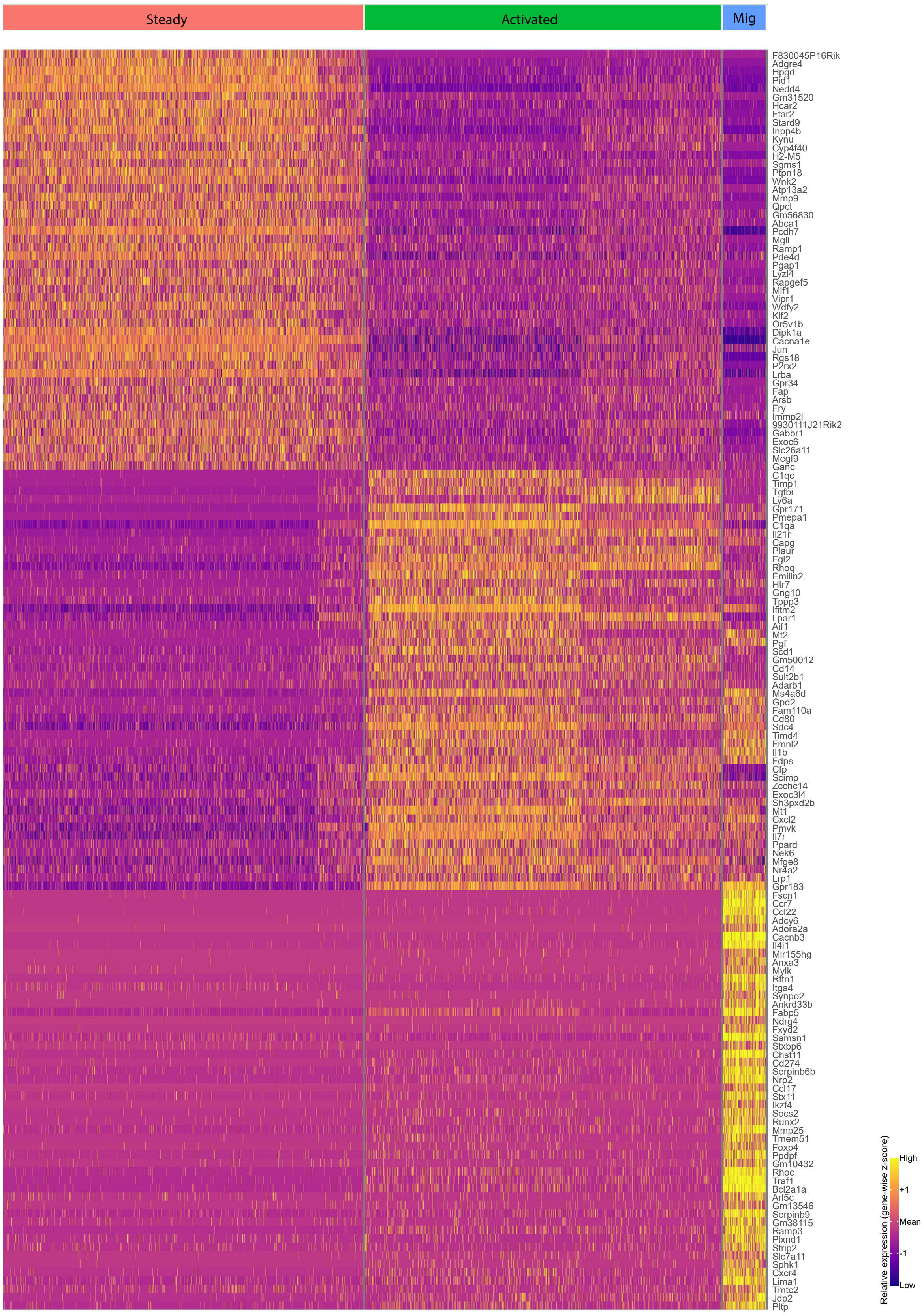
Steady
Activated
Mig
F830045P16Rik
Adgre4
Hpgd
Pid1
Nedd4
Gm31520
Hcar2
Ffar2
Stard9
Inpp4b
Kynu
Cyp4f40
H2-M5
Sgms1
Ptpn18
Wnk2
Atp13a2
Mmp9
Qpct
Gm56830
Abca1
Pcdh7
Mgll
Ramp1
Pde4d
Pgap1
Lyzl4
Rapgef5
Mlf1
Vipr1
Wdfy2
Klf2
Or5v1b
Dipk1a
Cacna1e
Jun
Rgs18
P2rx2
Lrba
Gpr34
Fap
Arsb
Fry
Immp2l
9930111J21Rik2
Gabbr1
Exoc6
Slc26a11
Megf9
Ganc
C1qc
Timp1
Tgfbi
Ly6a
Gpr171
Pmepa1
C1qa
Il21r
Capg
Plaur
Fgl2
Rhoq
Emilin2
Htr7
Gng10
Tppp3
Ifitm2
Lpar1
Aif1
Mt2
Pgf
Scd1
Gm50012
Cd14
Sult2b1
Adarb1
Ms4a6d
Gpd2
Fam110a
Cd80
Sdc4
Timd4
Fmnl2
Il1b
Fdps
Cfp
Scimp
Zcchc14
Exoc3l4
Sh3pxd2b
Mt1
Cxcl2
Pmvk
Il7r
Ppard
Nek6
Mfge8
Nr4a2
Lrp1
Gpr183
Fscn1
Ccr7
Ccl22
Adcy6
Adora2a
Cacnb3
Il4i1
Mir155hg
Anxa3
Mylk
Rftn1
Itga4
Synpo2
Ankrd33b
Fabp5
Ndrg4
Fxyd2
Samsn1
Stxbp6
Chst11
Cd274
Serpinb6b
Nrp2
Ccl17
Stx11
Ikzf4
Socs2
Runx2
Mmp25
Tmem51
Foxp4
Pdpdf
Gm10432
Rhoc
Traf1
Bcl2a1a
Arl5c
Gm13546
Serpinb9
Gm38115
Ramp3
Plxnd1
Strip2
Slc7a11
Sphk1
Cxcr4
Lima1
Tmtc2
Jdp2
Pltp
Relative expression (gene-wise z-score)
High
+1
Mean
-1
Low

### Figure S8

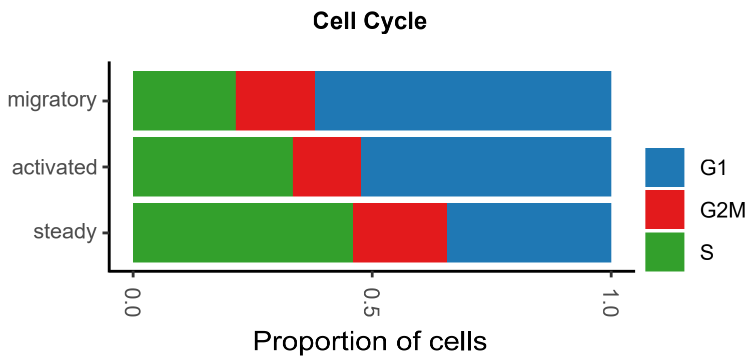
Cell Cycle
migratory
activated
steady
0.0
0.5
1.0
Proportion of cells
G1
G2M
S

### Figure S9

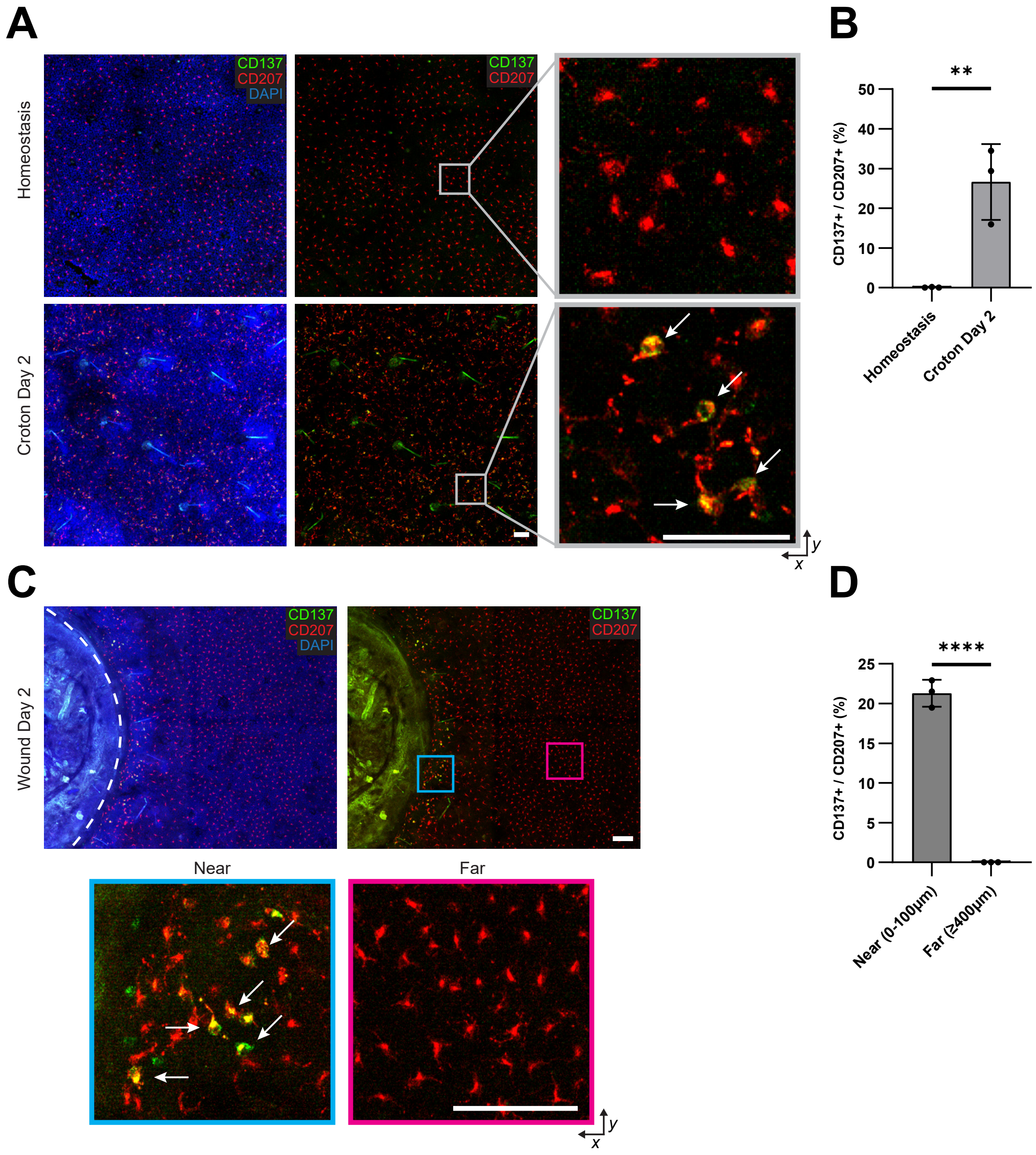

A
B
C
D
Homeostasis
Croton Day 2
Wound Day 2
CD137
CD207
DAPI
Near
Far
x
y
**
****
CD137+ / CD207+ (%)
0
10
20
30
40
50
5
15
25
Homeostasis
Croton Day 2
Near (0-100µm)
Far (≥400µm)

### Figure S10

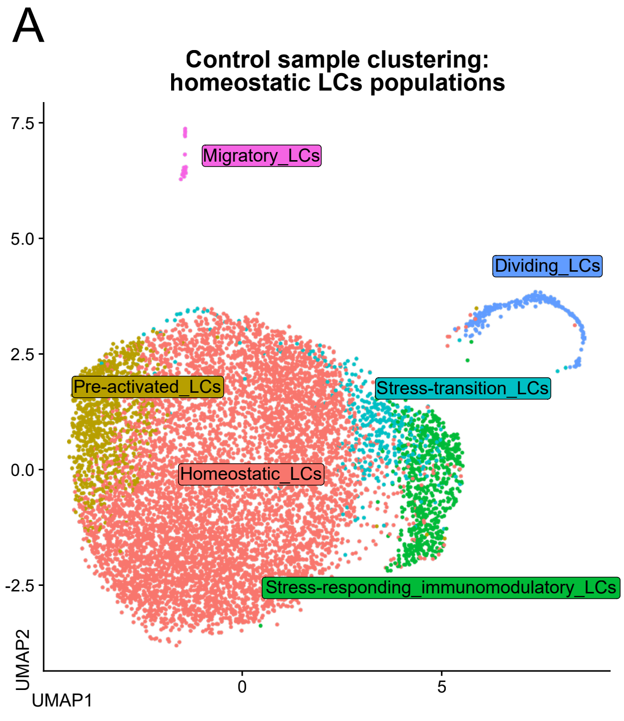
A
Control sample clustering:
homeostatic LCs populations
7.5
5.0
2.5
0.0
-2.5
Migratory_LCs
Dividing_LCs
Pre-activated_LCs
Stress-transition_LCs
Homeostatic_LCs
Stress-responding_immunomodulatory_LCs
UMAP2
UMAP1
0
5
B
CytoTRACE2 reveals a transcriptional program
consistent with a unipotent LC state
7.5
5.0
2.5
0.0
-2.5
Potency score
Totipotent
Pluripotent
Multipotent
Oligopotent
Unipotent
Differentiated
UMAP2
UMAP1
0
5

### Figure S11

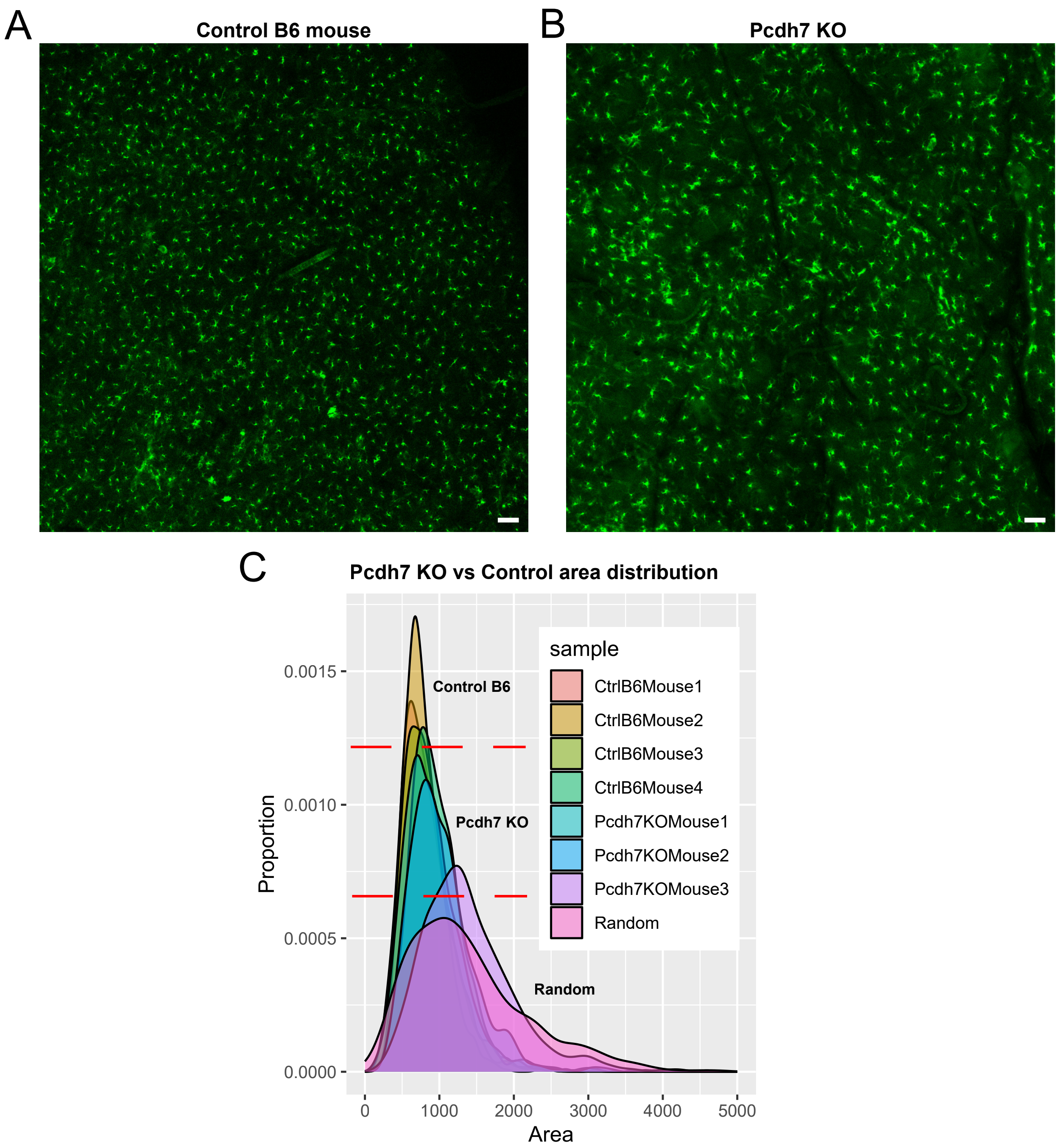
A
Control B6 mouse
B
Pcdh7 KO
C
Pcdh7 KO vs Control area distribution
Control B6
Pcdh7 KO
Random
sample
CtrlB6Mouse1
CtrlB6Mouse2
CtrlB6Mouse3
CtrlB6Mouse4
Pcdh7KOMouse1
Pcdh7KOMouse2
Pcdh7KOMouse3
Random
Proportion
0.0015
0.0010
0.0005
0.0000
0
1000
2000
3000
4000
5000
Area

### Figure S12

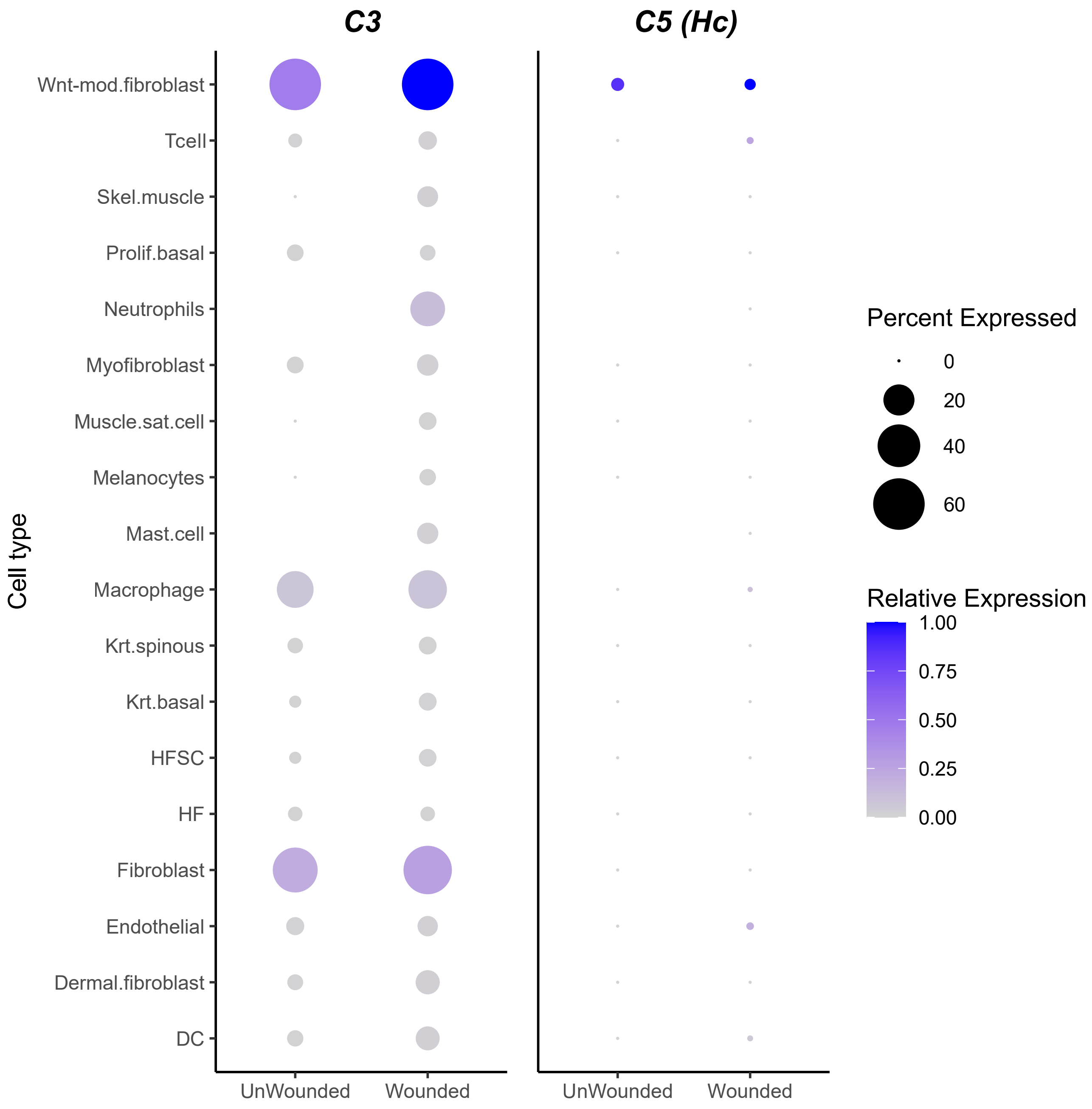

C3
C5 (Hc)
Wnt-mod.fibroblast
Tcell
Skel.muscle
Prolif.basal
Neutrophils
Myofibroblast
Muscle.sat.cell
Melanocytes
Mast.cell
Macrophage
Krt.spinous
Krt.basal
HFSC
HF
Fibroblast
Endothelial
Dermal.fibroblast
DC
Cell type
UnWounded
Wounded
UnWounded
Wounded
Percent Expressed
0
20
40
60
Relative Expression
1.00
0.75
0.50
0.25
0.00

### Figure S13

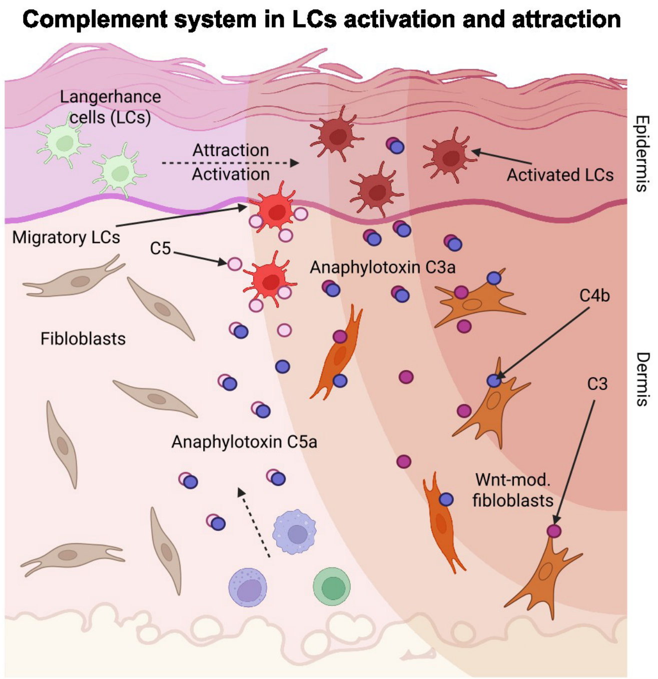
Complement system in LCs activation and attraction
Langerhance cells (LCs)
Attraction
Activation
Activated LCs
Epidermis
Migratory LCs
C5
Anaphylotoxin C3a
C4b
Fibloblasts
Dermis
C3
Anaphylotoxin C5a
Wnt-mod. fibloblasts
