## Supplemental Protocol 1 for "A Transcriptionally Distinct Intermediate Activation State Precedes Langerhans Cell Migration from the Epidermis"

### *Langerhans cells single-cell suspension preparation*

**Materials**:

- Sterile surgical scissors and forceps
- Nair cream (Naircare, USA)
- HBSS
- PBS
- FBS
- 70 μm cell strainer
- 60 mm plastic Petri dish
- 30 mm plastic Petri dish
- 50mL conical tubes
- 50 mg/ml Trypsin inhibitor (Sigma; T6522)
- 10 mg/ml DNaseI (Sigma; D4513)
- Dead Cell Removal Kit (Miltenyibiotec; 130-090-101)
- Epidermal Langerhans Cell MicroBead Kit, mouse (Miltenyibiotec; 130-095-408)
- LS columns (Miltenyibiotec, 130-042-401)
- QuadroMACS Separator (Miltenyibiotec, 130-091-051)
- Trypsin solution (0.3% trypsin, 150 mM NaCl, 0.5 mM KCl and 0.5 mM glucose)
- Single-cell suspension solution (HBSS, 5% FBS, 3ul 50 mg/ml Trypsin inhibitor, 3ul 10 mg/ml DNaseI)

**Note:**

- Hair should be removed as best as possible
- Avoid electric shaver
- If the experiment was initiated several days prior, reapply Nair cream to ensure complete hair removal before proceeding. Avoid re-nair of mechanically damaged conditions.

**Protocol:**

1. Euthanize mice, nair, cut off ears, keep in PBS.
2. Separate cartilage and epidermis-dermis sheet using forceps, keep in PBS at 37°C
3. Place the epidermis-dermis sheet, epidermis side up, onto the surface of the trypsin solution in a 60 mm Petri dish (up to 10 ears) and incubate for 1 hour and 45 minutes at 37°C.
    a. Prepare a "Petri grinder" by making 30-40 scratches along the inner wall of a 30 mm Petri dish using the blunt (non-sharp) edge of an inverted scalpel blade.
    b. Add 3 mL of single-cell suspension buffer to the prepared "Petri grinder."
4. Using two forceps, carefully separate the epidermis from the dermis. Gently rinse the epidermal sheet with PBS to remove residual debris and trypsin.
5. Using forceps, rub the epidermal sheet against the scratched surface of the "Petri grinder" to mechanically dissociate the cells.
6. Move the cloudy cell suspension in “Petri grinder” to a 37 °C and incubate for 15 minutes to enhance cell dissociation and inactivate trypsin
7. Pass the cell suspension through a 70 μm cell strainer into a clean 50ml tube. Rinse the strainer thoroughly with HBSS containing 5% FBS, bringing the final volume up to 50 mL.
8. Centrifuge the cell suspension at 300 × g for 20 minutes at 4 °C. A visible white cell pellet should form at the bottom of the tube.
9. Decant the HBSS from the tube in a single smooth motion, taking care not to disturb the cell pellet.
10. **Dead cell removing using Miltenyibiotec 130-090-101**
    a. Add 200 μL of Dead Cell Removal MicroBeads to the cell suspension and incubate for 15 minutes at room temperature
    b. Add 300ul LD Bind Buffer to the cell suspension
    c. Prepare LS column by adding 3 mL of LD Binding Buffer, place LS column on QuadroMACS Separator

d. Apply 500 µL of the cell suspension to the prepared LS column. Collect the flow-through into a 50 mL tube. Rinse the column four times with 3 mL of LD Binding Buffer, pool all flow-through fractions in 50 ml tube (do not remove LS column from separator at any step).

**11. Langerhans cells isolation using Miltenyibiotec 130-095-408**

a. Centrifuge the cell suspension at 300 × g for 20 minutes at 4 °C. A visible white cell pellet should form at the bottom of the tube.
b. Decant the LD Binding Buffer from the tube in a single smooth motion, taking care not to disturb the cell pellet.
c. Add 80ul AB Buffer EDTA to cell suspension
d. Add 10ul FC block solution to cell suspension incubate 10 minutes at 8 °C
e. Add 15ul Cd207 beads to cell suspension incubate 15 minutes at 8 °C
f. Add 1 ml AB Buffer EDTA, centrifuge at 300 × g for 10 minutes at 4 °C
g. Prepare LS column by adding 3 mL of AB Buffer EDTA, place LS column on QuadroMACS Separator
h. Resuspend cells-Cd207 beads suspension in 1 ml AB Buffer EDTA
i. Apply 1 ml of the cell suspension to the prepared LS column. Rinse the column four times with 3 mL of AB Buffer EDTA. Discard the flow-through.
j. Add 5 ml AB Buffer NO EDTA to column, remove, from QuadroMACS Separator
k. Using the plunger, gently push the retained cells from the first LS column onto a new, pre-equilibrated LS column.
l. Rinse the column four times with 3 mL of AB Buffer NO EDTA. Discard the flow-through.
m. Add 5 ml AB Buffer NO EDTA to column, remove, from QuadroMACS Separator
n. Using the plunger, gently push the retained cells from the second LS column onto 50 ml tube.
o. Centrifuge the cell suspension at 300 × g for 20 minutes at 4 °C, cell pellet should be visible.
p. Resuspend in the necessary volume of AB Buffer NO EDTA
q. Pass the cell suspension through a 30 µm cell strainer
r. Procced standard 10x library preparation QC steps (FACS, Trypan Blue)
